# Targeted DNA integration in human cells without double-strand breaks using CRISPR RNA-guided transposases

**DOI:** 10.1101/2023.03.17.533036

**Authors:** George D. Lampe, Rebeca T. King, Tyler S. Halpin-Healy, Sanne E. Klompe, Marcus I. Hogan, Phuc Leo H. Vo, Stephen Tang, Alejandro Chavez, Samuel H. Sternberg

## Abstract

Traditional genome-editing reagents such as CRISPR-Cas9 achieve targeted DNA modification by introducing double-strand breaks (DSBs), thereby stimulating localized DNA repair by endogenous cellular repair factors. While highly effective at generating heterogenous knockout mutations, this approach suffers from undesirable byproducts and an inability to control product purity. Here we develop a system in human cells for programmable, DSB-free DNA integration using Type I CRISPR-associated transposons (CASTs). To adapt our previously described CAST systems, we optimized DNA targeting by the QCascade complex through a comprehensive assessment of protein design, and we developed potent transcriptional activators by exploiting the multi-valent recruitment of the AAA+ ATPase, TnsC, to genomic sites targeted by QCascade. After initial detection of plasmid-based transposition, we screened 15 homologous CAST systems from a wide range of bacterial hosts, identified a CAST homolog from *Pseudoalteromonas* that exhibited improved activity, and increased integration efficiencies through parameter optimization. We further discovered that bacterial ClpX enhances genomic integration by multiple orders of magnitude, and we propose that this critical accessory factor functions to drive active disassembly of the post-transposition CAST complex, akin to its demonstrated role in Mu transposition. Our work highlights the ability to functionally reconstitute complex, multi-component machineries in human cells, and establishes a strong foundation to realize the full potential of CRISPR-associated transposons for human genome engineering.

---

Since the pioneering work of Jasin and colleagues established that targeted DNA double-strand breaks (DSBs) drastically increase rates of recombination^1, 2^, genome editing technologies have primarily focused on discovering and developing nucleases with user-defined specificities^3^. RNA-guided DNA endonucleases encoded by CRISPR-Cas systems, including Cas9 and Cas12, offer ease of programmability and high-efficiency activity in a wide range of cells and organisms, and have therefore experienced widespread adoption for basic research, agricultural applications, and human therapeutics^4, 5^. In mammalian cells, Cas9-mediated DSBs are primarily repaired in one of two ways — non-homologous end joining (NHEJ) and homology-directed repair (HDR) — with the efficiency of NHEJ typically exceeding that of HDR by at least an order of magnitude^6^. Although improved methods of HDR-based insertion have begun to emerge^7–10^, precise modifications necessitating larger cargo sizes remain inefficient and difficult to generate, particularly in cell types that do not express sufficient levels of recombination machinery^11–13^.

Recent studies have further highlighted the range of undesirable (and previously undetected) byproducts of DSB-based genome editing, including large-scale genomic deletions, chromosomal translocations, and chromothripsis^14–17^, which confound results and pose serious safety concerns. Additionally, researchers have shown that *p53*-mediated DNA damage responses are induced by Cas9-generated DSBs, suggesting that genome editing techniques may select for *p53*-mutated, potentially tumorigenic cells^18, 19^. “Second-generation” editing reagents, including base editors and prime editors, exploit CRISPR-Cas9 for programmable RNA-guided DNA targeting while leveraging fused effector domains to perform site-specific chemistry on the genome, enabling precise, DSB-independent modifications^20–22^. However, both approaches have traditionally been restricted to edits ranging from single to ∼50 base pairs (bp), rendering insertion of larger payloads inaccessible. Novel “third-generation” editing reagents combining prime editors with serine integrases, such as TwinPE and PASTE, have been shown to enable larger DNA insertions and other modifications; however these methods still rely on previously mentioned complex editing methods requiring resolution of DNA intermediates that generate additional undesired indels at the target site and incomplete modifications when the multiple necessary enzymatic events do not occur in concert^23, 24^.

Numerous biotechnology applications rely on genomic insertion of kilobase-scale genetic cassettes, including conventional gene therapy, crop engineering, biologics production, and metabolic engineering^25–27^. Lentiviral vectors are a highly utilized gene delivery vehicle because they integrate with high efficiency across diverse cell types, but they exhibit promiscuous specificity, offer little control over copy number, present limitations in cargo capacity and design, and require numerous manufacturing steps^28^. Transposases such as Sleeping Beauty and piggyBac also integrate DNA without relying on host recombination and can better accommodate large payloads, but they lack specificity and copy number control^29–31^. In contrast to these approaches, recombinases such as Cre and Bxb1 offer excellent specificity and product purity, but are not programmable and thus require researchers to first generate engineered cells containing the obligate recombination site^32, 33^. The ideal DNA integration technology would function in a single step and avoid generating DSBs or indels, while retaining the programmability afforded by RNA-guided DNA targeting.

Recent studies have attempted to engineer RNA-guided transposases by fusing Cas9 to various transposase domains, but these efforts have remained reliant on DSBs and/or failed to achieve stringent specificity control^34–37^. In contrast, bacterial CRISPR-associated transposases (CASTs) catalyze insertion of large DNA payloads in a targeted manner without DSBs. Using a Type I-F CAST system derived from *Vibrio cholerae* Tn*6677*, we recently reported DSB-free DNA insertions in multiple bacterial species and demonstrated that this approach exhibited exquisite genome-wide specificity and could be easily reprogrammed to user-defined sites with single-bp accuracy^38, 39^. Long-read whole-genome sequencing confirmed the purity of integration products, and additional heterologous reconstitution experiments demonstrated enzymatic function independent of obligate recombination factors^39, 40^. We therefore sought to leverage RNA-guided transposases for targeted DNA integration in mammalian cells, despite the formidable obstacle of reconstituting a complex, multi-component pathway that depends on a donor DNA, guide CRISPR RNA (crRNA), and assembly of seven distinct proteins, many of which function in an oligomeric state (**Fig. 1a,b**).

**Fig. 1.**
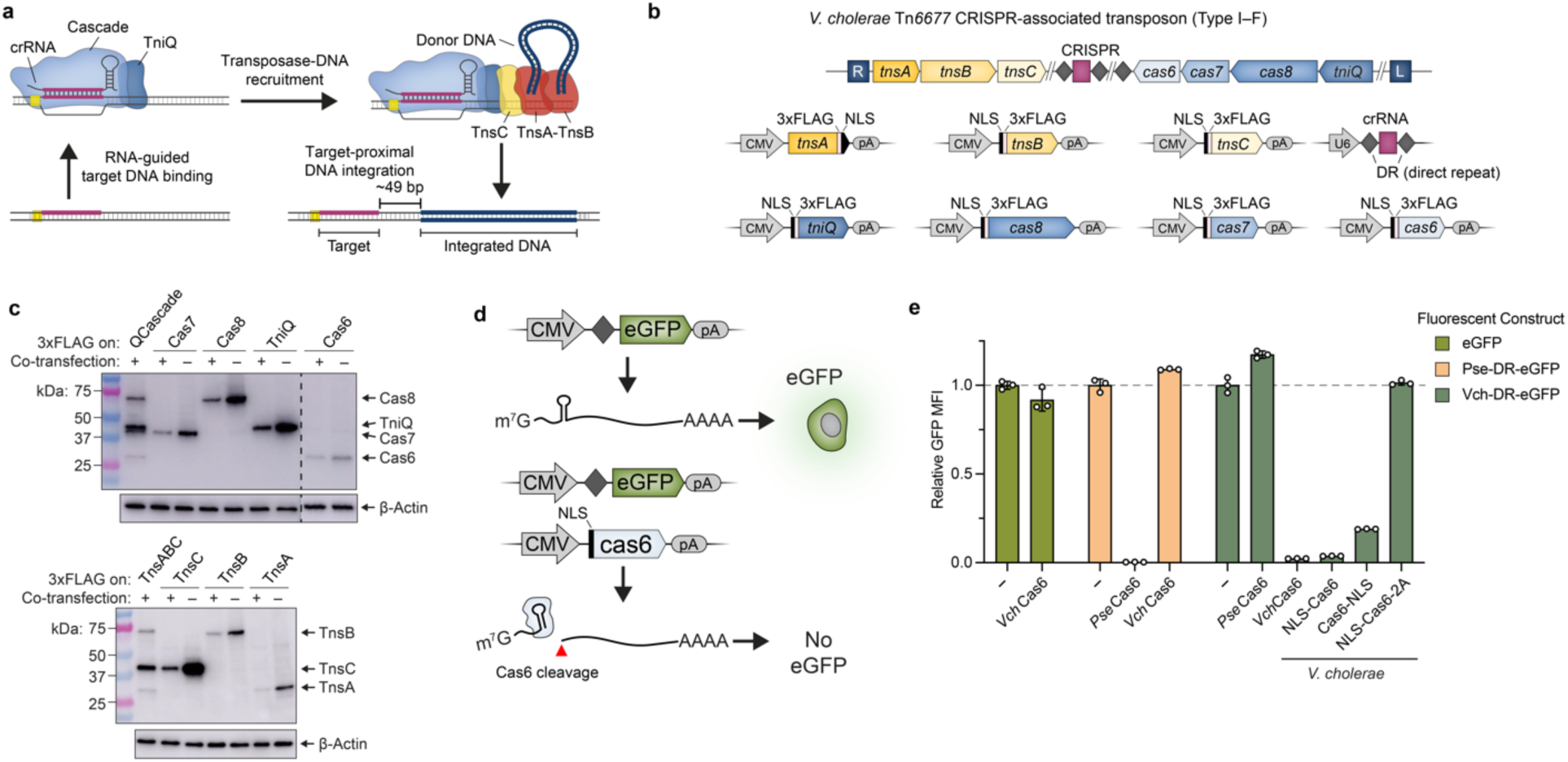
Reconstitution of protein-RNA CAST components in human cells. **a,** Schematic detailing DNA integration using RNA-guided transposases. **b,** Type I-F CRISPR-associated transposons encode the CRISPR RNA and seven proteins needed for DNA integration (top). Mammalian expression vectors used for heterologous reconstitution in human cells are shown at bottom. **c,** Western blotting with anti-FLAG antibody demonstrates robust protein expression upon individual (–) or multi-plasmid (+) co-transfection of HEK293T cells. Co-transfections contained all *Vch*CAST components, with the FLAG-tagged subunit(s) indicated. β-actin was used as a loading control. **d,** Schematic of eGFP knockdown assay to monitor crRNA processing by Cas6 in HEK293T cells. Cleavage of the CRISPR direct repeat (DR)-encoded stem-loop severs the 5′-cap from the ORF and polyA (pA) tail, leading to a loss of eGFP fluorescence (bottom). **e,** Transposon-encoded *Vch*Cas6 (Type I-F3) exhibits efficient RNA cleavage and eGFP knockdown, as measured by flow cytometry. Knockdown was comparable to *Pse*Cas6 from a canonical CRISPR-Cas system (Type I-E)^40^, was absent with a non-cognate DR substrate, and was sensitive to C-terminal tagging. To control for over-expression, data were normalized to negative control conditions (–), in which dCas9 was co-transfected with the reporter. Data are shown as mean ± s.d. for n = 3 biologically independent samples.

Here we report mammalian CAST activity using two diverse systems from *V. cholerae* and *Pseudoalteromonas*^41^, demonstrating that the same molecular determinants of RNA-guided transposition hold true in bacteria and eukaryotes. Intriguingly, integration efficiencies were initially much lower at endogenous target sites compared to episomal plasmid substrates, which led us to identify bacterial ClpX as a critical accessory factor that enhanced genomic integration by more than two orders of magnitude. During our engineering efforts, we also developed a strategy for targeted recruitment of an oligomeric transposase component, TnsC, which we harnessed to achieve potent transcriptional activation at levels similar to conventional dCas9-based reagents. Taken together with recent studies harnessing alternative Type I CRISPR-Cas systems for eukaryotic genome and transcriptome engineering^42–46^, our work challenges the reliance on single-effector editing reagents and provides a strong starting point for genome engineering using RNA-guided, CRISPR-associated transposases.

## RESULTS

### Heterologous expression of CAST components in human cells

Bacterial Tn7-like transposons have co-opted at least three distinct types of nuclease-deficient CRISPR-Cas systems for RNA-guided transposition (I-B, I-F, and V-K)^36, 45, 46^, with each exhibiting unique features. We carefully reviewed fidelity and programmability parameters for experimentally characterized CAST systems, alongside recently described Cas9-transposase fusion approaches^32–35^, and opted to focus our efforts on the Type I-F *V. cholerae* CAST (*Vch*CAST; previously also referred to as *Vch*INTEGRATE) because of its optimal integration efficiency, specificity, and absence of cointegrates^38–40, 47^. Within this system, a ribonucleoprotein complex comprising TniQ and Cascade (*Vch*QCascade) performs RNA-guided DNA targeting, thereby defining sites for transposon DNA insertion^36, 48^. Excision and integration reactions are catalyzed by the heteromeric TnsA-TnsB transposase, but only after prior recruitment of the AAA+ ATPase, TnsC^48, 49^. Integrated DNA payloads must be flanked by transposon left and right end sequences, which encode TnsB binding sites and define boundaries of the mobile element.

We adopted a methodical, bottom-up approach to port *Vch*CAST into human cells. To first establish whether the component parts were efficiently expressed, we cloned each protein-coding gene onto a standard mammalian expression vector with an N- or C-terminal nuclear localization signal (NLS) and 3xFLAG epitope tag (**Fig. 1b**). Using Western blotting, we showed robust heterologous protein expression, both individually and when all CAST proteins were co-expressed (**Fig. 1c**). Cellular fractionation provided evidence of nuclear trafficking, and we also demonstrated efficient expression and trafficking of an engineered TnsAB fusion protein (TnsAB_f_) that we previously showed retains wild-type activity (**Supplementary Fig. 1**)^40^. However, initial attempts to reconstitute RNA-guided DNA integration in HEK293T cells proved unsuccessful, even after exploring numerous strategies to enrich rare events through both positive and negative selection (data not shown). We therefore decided to separately assess guide RNA expression by adapting a previously developed approach^43^ to monitor crRNA biogenesis within the 5′ untranslated region (UTR) of a GFP-encoding mRNA. Cas6 is a ribonuclease subunit of Cascade that cleaves the CRISPR repeat sequence in most Type I CRISPR-Cas systems^50^, which in our assay would sever the 5′ cap from the GFP open reading frame and thus lead to fluorescence knockdown (**Fig. 1d**). Accordingly, we observed near-total loss of GFP fluorescence when the reporter plasmid was co-transfected with cognate *Vch*Cas6, but not when the reporter encoded a non-cognate CRISPR repeat or lacked a repeat altogether (**Fig. 1e**). Interestingly, GFP knockdown was substantially reduced when Cas6 contained a C-terminal NLS or 2A peptide (**Fig. 1e**), indicating a sensitivity to terminal tagging that could not be easily explained by the cryoEM structure (see below)^51^. Collectively, these experiments verified expression of all protein and RNA components from *Vch*CAST, leading us to next focus on functional reconstitution of RNA-guided DNA targeting by QCascade.

### QCascade and TnsC function as transcriptional activators

Unlike most Type II and V CRISPR-Cas systems, which encode single-effector proteins that function as RNA-guided DNA nucleases (Cas9 and Cas12, respectively), the Cascade complex encoded by Type I systems does not possess DNA cleavage activity and instead exhibits long-lived target DNA binding upon R-loop formation, analogously to catalytically inactive Cas9 (dCas9)^52^. We decided to leverage this activity for transcriptional activation of an mCherry reporter gene by fusing transcriptional activators to QCascade, as recently done for other Type I systems^43, 44, 46^, thereby converting DNA binding into a detectable signal that would allow facile troubleshooting and optimization of QCascade function (**Supplementary Fig. 2a**).

We first constructed activators using a Type I-E Cascade unrelated to transposons from *Pseudomonas* sp. S-6–2 (*Pse*Cascade_IE), which we previously exploited for genome engineering in human cells^42^. We fused VP64 to the hexameric Cas7 subunit and concatenated all five *cas* genes within a single polycistronic vector downstream of a CMV promoter, by linking them together with virally derived 2A ‘skipping’ peptides; the crRNA was separately expressed from a U6 promoter (**Supplementary Fig. 2a**). The resulting expression plasmids yielded ∼260-fold mCherry activation when co-transfected with the reporter plasmid, similar to levels achieved with dCas9-VPR, and the effect was ablated in the presence of a non-targeting crRNA (**Fig. 2b**). Surprisingly, when we tested nearly identical designs using the transposon-encoded Type I-F QCascade homolog from *V. cholerae*, we failed to detect any activation (**Fig. 2a,b**).

**Fig. 2.**
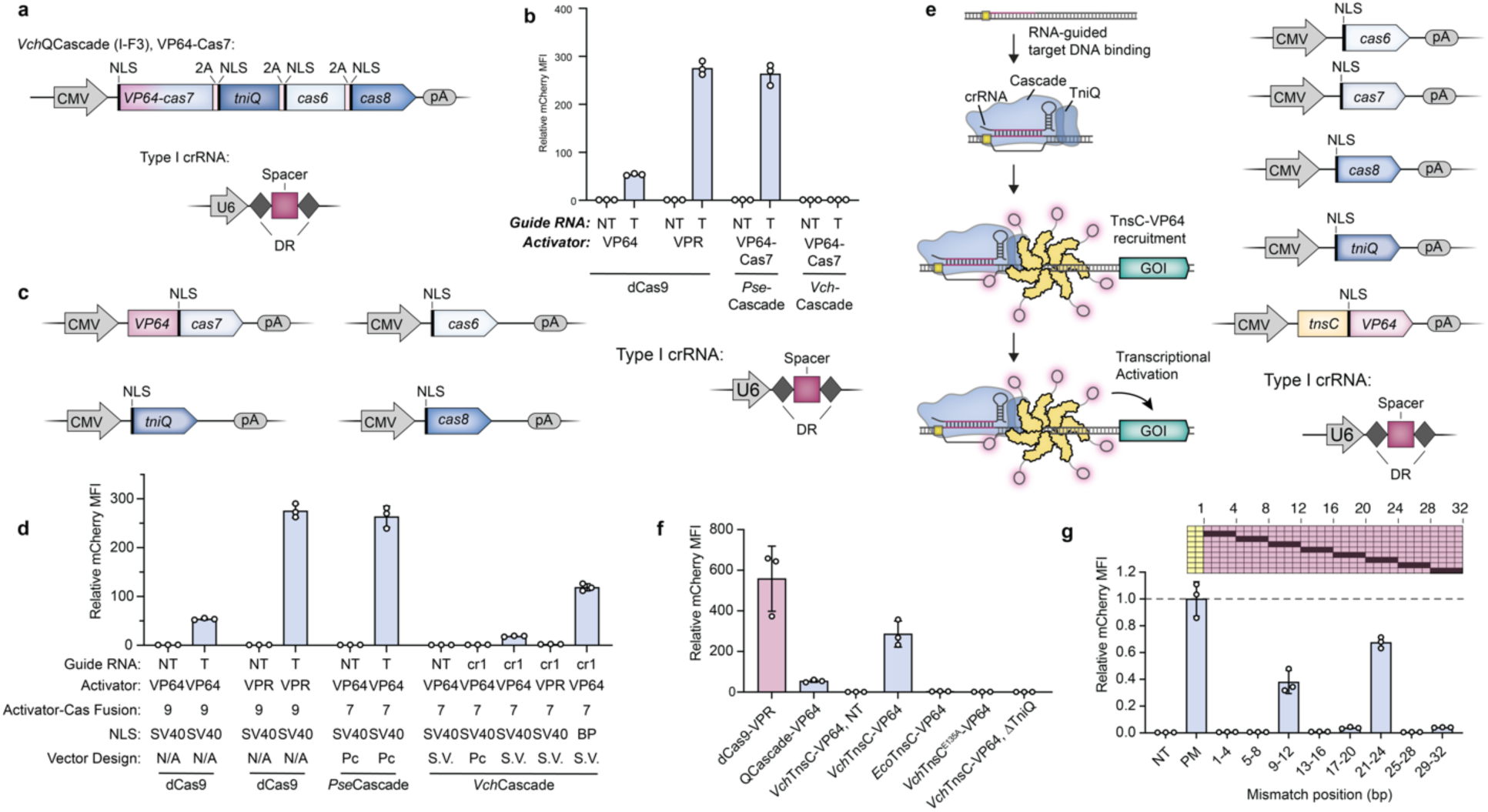
Development of QCascade and TnsC-based transcriptional activators to monitor DNA targeting. **a,** Design of mammalian expression vectors encoding transposon-encoded Type I-F3 systems (*Vch*QCascade). Cascade subunits are concatenated on a single polycistronic vector and connected by virally derived 2A peptides, as described previously^40^. **b,** Normalized mCherry fluorescence levels for the indicated experimental conditions, measured by flow cytometry. Whereas *Pse*Cascade stimulated robust activation, *Vch*QCascade was inactive under these conditions. NT, non-targeting sgRNA/crRNA; T, targeting sgRNA/crRNA. **c,** Design of separately encoded *Vch*QCascade mammalian expression vectors with optimized NLS tag placement. **d,** *Vch*QCascade mediates transcriptional activation when encoded by re-engineered expression vectors, as measured by flow cytometry. mCherry expression is further enhanced when replacing mono-partite (SV40) NLS tags with bipartite (BP) NLS tags. NT, non-targeting; T, targeting. **e,** Schematic of transcriptional activation assay, in which DNA targeting by *Vch*QCascade leads to multi-valent recruitment of *Vch*TnsC-VP64. The assembly mechanism is based on our recent biochemical, structural, and functional data^60^. **f,** Normalized mCherry fluorescence levels for the indicated experimental conditions, measured by flow cytometry. *Vch*TnsC-based activation requires cognate protein-protein interactions, is strictly dependent on the presence of TniQ, and involves ATP-dependent oligomer formation, which is eliminated with the E135A mutation. Several controls are shown for comparison, and guide RNAs target the same sites shown in **Supplementary Figure 3a**. NT, non-targeting crRNA. **g,** Transcriptional activation shows strong sensitivity to RNA-DNA mismatches within both the PAM-proximal seed sequence and a PAM-distal region implicated in TnsC recruitment. Data are shown as in **f**, and the schematic at top displays the mismatched positions that were tested. Data were normalized to the perfectly matching (PM) crRNA. Data in **b, d, f, g** are shown as mean ± s.d. for n = 3 biologically independent samples.

We suspected that the presence of N-terminal NLS tags, C-terminal 2A tags, or both, might be inhibiting QCascade assembly and/or RNA-guided DNA targeting, despite the fact that all termini appeared to be solvent-accessible in our experimentally determined *Vch*QCascade structure (**Supplementary Fig. 2b**)^51^. To systematically investigate this possibility, we cloned peptide tags onto the termini of all *Vch*CAST components and tested their impact in *E. coli* transposition assays. While some tags had little effect on activity, others led to a severe reduction or complete loss of targeted DNA integration (**Supplementary Fig. 2c**), highlighting the sensitivity of this system to minor perturbations. The transposase components were particularly vulnerable, with an N-terminal tag on TnsA and C-terminal tags on TnsB and TnsC being largely prohibitive. Within the context of QCascade, C-terminal 2A tags on TniQ and Cas7 each reduced integration by >90%, which could explain the lack of transcriptional activation we observed using polycistronic vector designs. We also screened multiple components for activator fusions and found that the N-terminus of Cas7 was amenable to both VP64 and VPR fusions in bacteria (**Supplementary Fig. 2d**).

With these data in hand, we retested QCascade-VP64 in human cells using individual expression vectors with optimized NLS tag locations for each component, and detected mCherry activation for two distinct crRNAs, evidencing successful assembly and target binding in human cells (**Fig. 2c,d Supplementary Fig 2e**). Activation levels were further increased by replacing all monopartite SV40 NLS tags with bipartite (BP) NLS tags, and this activity was strictly dependent on the simultaneous expression of Cas8, Cas7, Cas6, and a targeting crRNA (**Fig. 2d, Supplementary Fig. 2e,f**). Interestingly, although Cas7 tolerated a VPR fusion in bacteria, we were unable to detect transcriptional activation in mammalian cells using VPR-Cas7 (**Fig. 2d, Supplementary Fig. 2d,e**). These results highlighted the importance of carefully dissecting the effects of all sequence modifications being introduced to *Vch*CAST components, even those appearing innocuous, and emphasized the value of fluorescence reporter assays in debugging molecular events upstream of DNA integration.

Early dCas9-based transcriptional activators revolved around recruitment of an activator domain covalently linked to a single dCas9^53–55^, whereas later methods have exploited strategies for multivalent recruitment of one or more effector domains^56, 57^. In the case of CAST systems, recent experiments have demonstrated that TnsC forms large ATP-dependent oligomers that assemble onto double-stranded DNA (dsDNA) and are specifically recruited to DNA-bound QCascade with high genome-wide specificity in *E. coli*^49, 58, 59^. We thus hypothesized that these properties could be leveraged for multivalent assembly of TnsC to increase the potency of transcriptional activation in mammalian cells, while also demonstrating recruitment of a critical transposase component in a QCascade-dependent fashion (**Fig. 2e**).

We fused VP64 to either the N- or C-terminus of TnsC, targeted seven candidate sites upstream of our mCherry reporter gene (**Supplementary Fig. 3a**), and investigated the potential for TnsC to stimulate transcriptional activation. Strikingly, TnsC-VP64 activators drove substantially higher levels of mCherry activation than QCascade alone, and activation levels could be further improved by optimizing the relative amount of each expression plasmid used during transfection (**Fig. 2f, Supplementary Fig. 3b**). This effect was absent when TniQ was omitted or an *E. coli* TnsC homolog was substituted, confirming the importance of cognate TniQ-TnsC interactions. Furthermore, a TnsC ATPase mutant that prevents oligomer formation (E135A)^49^ also abolished transcriptional activation, suggesting that the observed signal requires protein oligomerization on DNA (**Fig. 2f**). TnsC homologs from Type V-K CAST systems form filaments non-specifically on dsDNA^58, 59^, and we were therefore keen to investigate the fidelity of *Vch*TnsC- mediated activation. Non-targeting controls generated undetectable mCherry MFI above background levels, demonstrating the specificity of potential TnsC filamentation in Type I-F CASTs (**Fig. 2f)**. When probing the specificity of QCascade DNA binding, intermediate levels of transcriptional activation were retained when mismatches were tiled within the middle of the 32- bp target site, but there was a strict requirement for cognate pairing in the seed (positions 1–8) and PAM-distal (positions 25–32) regions (**Fig. 2g**).

Having demonstrated the ability of TnsC-based activation to potently induce expression of a reporter gene, we targeted four endogenous genes in the human genome (*TTN*, *MIAT*, *ASCL1*, and *ACTC1*), which have been previously targeted with CRISPRa using dCas9-VPR^60^. We designed three or four distinct crRNAs tiled upstream of the transcription start site and delivered them by either transfecting a single crRNA expression plasmid, co-transfecting multiple crRNA expression plasmids, or transfecting a single crRNA expression plasmid containing a four-spacer CRISPR array (**Fig. 3a, Supplementary Fig. 3c,d**). *TTN* induction by TnsC-VP64 was comparable to dCas9-VP64 and dCas9-VPR activation, and consistent with our model, the presence of Cas8 and TniQ were strictly required (**Fig. 3a**). Potent activation was seen on other genomic targets ranging from 200-fold (*MIAT*) to >1000-fold (*ASCL1*), highlighting the programmability of our multimeric system (**Fig. 3a**), though other sites showed more moderate activation (**Supplementary Fig. 3e**). Furthermore, we demonstrated the ability to utilize a multiplexed CRISPR array containing four spacers that each targeted a different gene to achieve robust transcriptional activation of all 4 genes (*TTN*, *MIAT*, *ASCL1*, and *ACTC1*) in the same cell population at levels comparable to activation achieved by single spacer CRISPR arrays (**Fig. 3b, 3c)**.

**Fig. 3.**
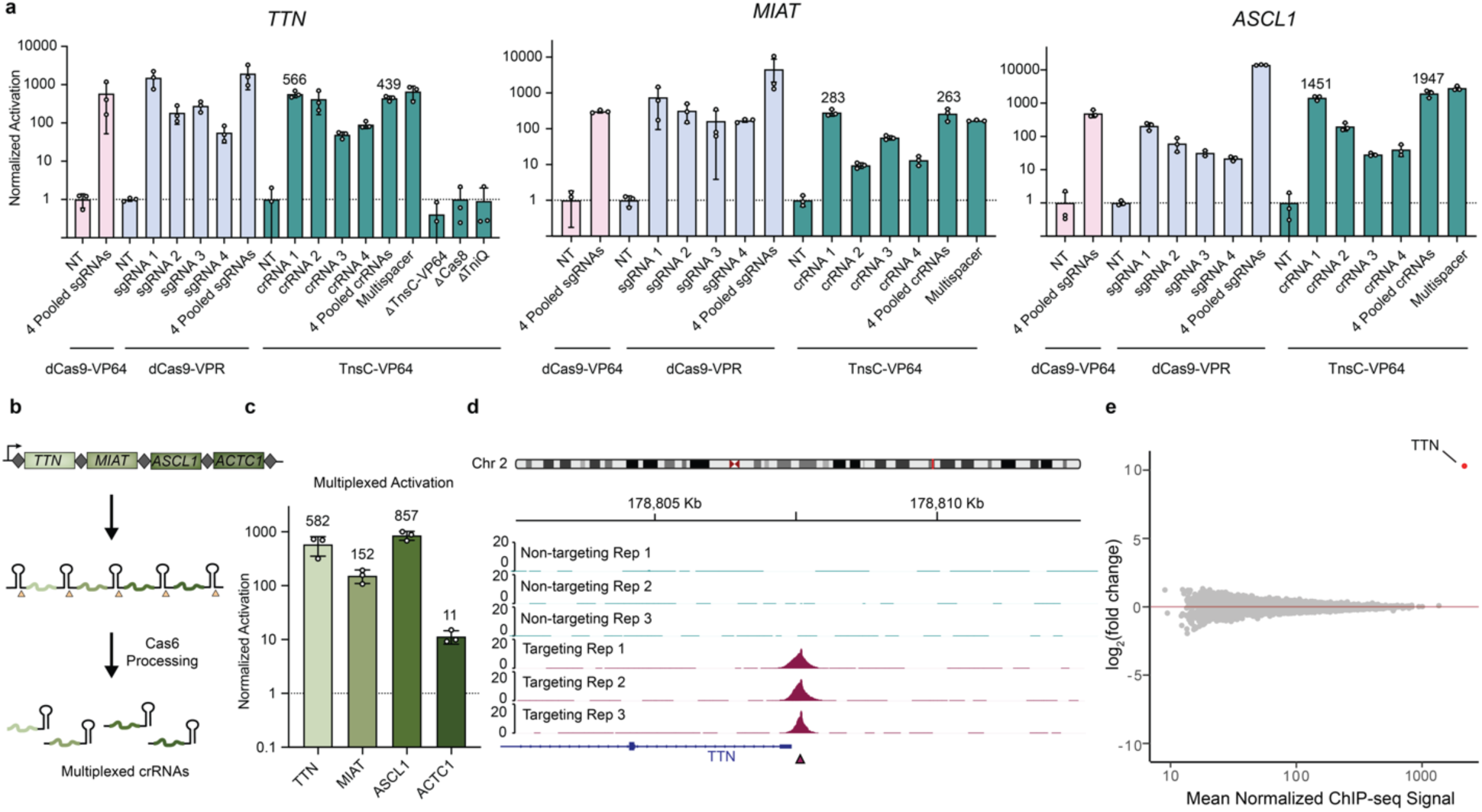
Potent genomic transcriptional activation via RNA-guided recruitment of the AAA+ ATPase, TnsC. **a,** TnsC-VP64 directs efficient transcriptional activation of endogenous human gene expression, as measured by RT-qPCR. Four distinct crRNAs were combined for each condition and were either delivered individually, as a pool, or as a single multi-spacer multiplexed CRISPR array. The dCas9-VP64 and dCas9-VPR comparisons utilized four distinct sgRNAs encoded on separate plasmids. NT, non-targeting; T, targeting. **b,** Schematic demonstrating Cas6’s ability to process CRISPR arrays in vivo, thus allowing for the use of multiplexed CRISPR arrays to target multiple sites concurrently. **c,** Multiplexed activation of 4 distinct genes in the same cell pool. **d,** 10 kb viewing window of ChIP-seq signal at the *TTN* promoter corresponding to *TTN* Guide 1. **e,** Differential binding analysis plot. Across consensus peaks for each condition, the only region exhibiting significantly different ChIP enrichment (FDR < 0.05) between targeting and non-targeting conditions was the peak at the *TTN* promoter. Data in **a,c** are shown as mean ± s.d. for n = 3 biologically independent samples. Viewing windows in **d,** are shown for 3 biologically independent targeting and non-targeting samples. Data in **e,** is shown as the mean for n = 3 biologically independent samples for each condition on the y axis, and the mean for all n = 6 biologically independent samples on the x axis, irrespective of condition.

We next investigated the fidelity of TnsC recruitment by performing ChIP-seq after co-transfecting plasmids encoding FLAG-tagged TnsC, protein components of QCascade, and a *TTN*- specific crRNA. Analysis of the resulting data revealed a sharp peak directly upstream of the *TTN* transcriptional start site (TSS) at the expected target site, which was absent in non-targeting (NT) samples transfected with a crRNA containing a spacer not found in the human genome (**Fig. 3d, Supplementary Fig. 4a,b)**. To assess off-target binding, we analyzed all peaks in both targeting and non-targeting conditions across three biological replicates and performed differential binding analysis, revealing only a single region at the *TTN* promoter that exhibited significantly different binding affinity between both conditions (FDR < 0.05)^61^, highlighting the specificity of Type I-F CAST assembly (**Supplementary Fig. 4c, Fig. 3e**). Heatmap analysis of additional peaks that were called in either targeting or non-targeting conditions revealed low enrichment values, and a further manual inspection of 5 potential off-target sites that exhibited high similarity to the *TTN* spacer sequence lacked any detectable signal enrichment in the ChIP-seq datasets (**Supplementary Fig. 4d-g**). Together with our recent study of *Vch*CAST factor recruitment in *E. coli*^49^, these results indicate that TnsC binds target sites marked by QCascade with high-fidelity, and that the intrinsic ability of TnsC to form ATP-dependent oligomers enables multiple copies of an effector protein to be delivered to genomic sites targeted by a single guide RNA.

This programmable, multivalent recruitment represents an exciting opportunity to further develop genome and transcriptome engineering tools that benefit from RNA-guided DNA binding of an effector ATPase. In the context of efforts to reconstitute CAST systems, TnsC-mediated transcriptional activation provided compelling evidence that both CRISPR- and transposon-associated protein components can be functionally assembled at plasmid and genomic target sites in a highly specific and programmable manner, encouraging our efforts to next probe for RNA-guided DNA integration.

### RNA-guided episomal DNA integration in human cells

We reasoned that the baseline efficiency of RNA-guided transposition might be low prior to optimization, and therefore sought to develop a sensitive assay that would enrich integration products. We cloned a promoter-driven chloramphenicol resistance cassette (CmR) within the mini-transposon of a donor plasmid (pDonor) and then targeted the same sequence on the mCherry reporter plasmid (pTarget) that was used in transcriptional activation experiments. Upon successful transposition in HEK293T cells, integrated pTarget products will carry both CmR and KanR drug markers and can thus be selected for by transforming *E. coli* with plasmid DNA isolated from transfected cells (**Fig. 4a**). Importantly, in these experiments we used a pDonor backbone that cannot be replicated in standard *E. coli* strains, reducing background from unreacted plasmids. We also opted to use a TnsAB fusion protein (TnsAB_f_)^40^ that contains an internal bipartite NLS and maintains wild-type activity in *E. coli* (**Supplementary Fig. 1c**), thereby reducing the number of unique protein components.

**Fig. 4.**
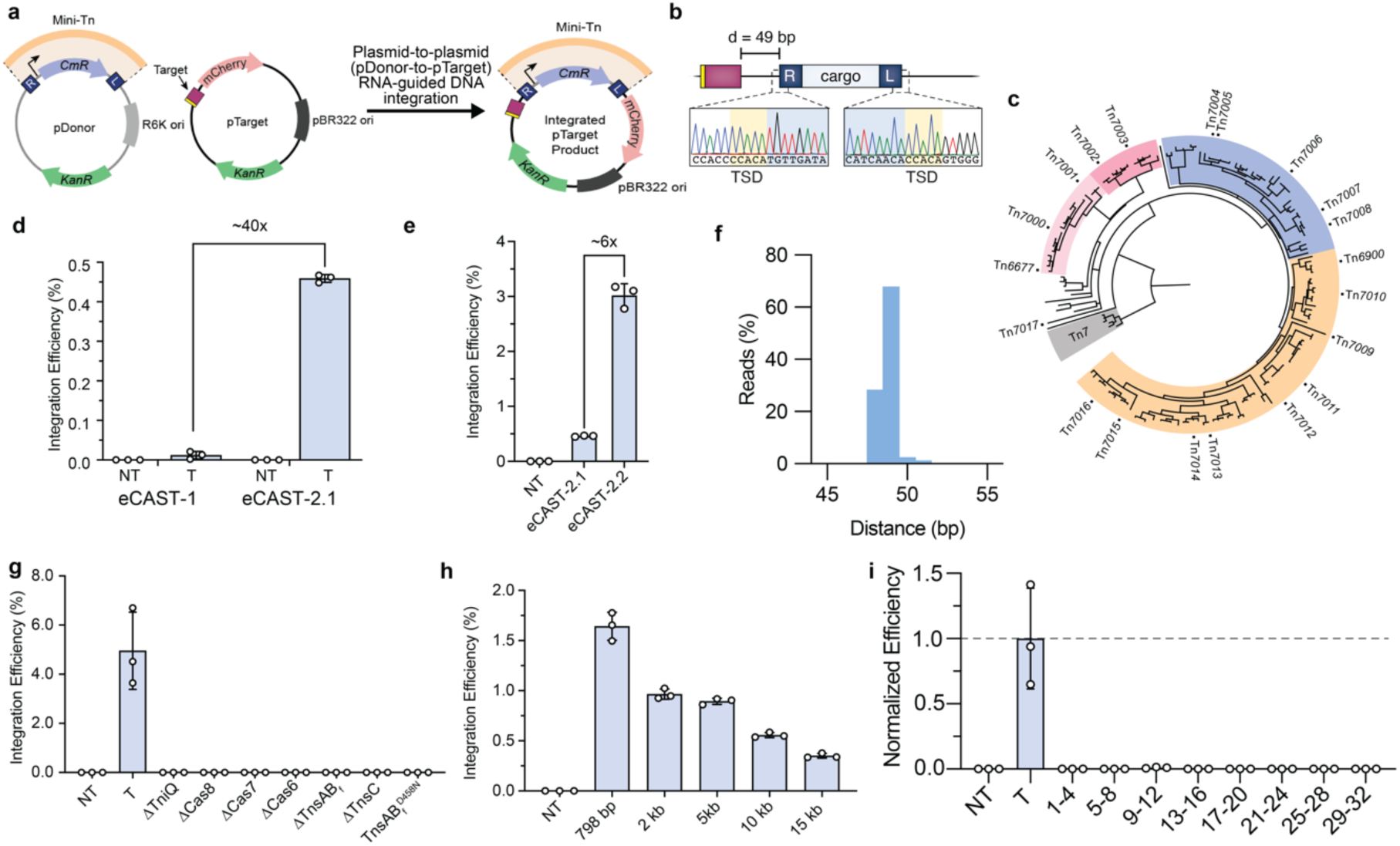
Plasmid-based RNA-guided DNA integration in human cells using diverse CRISPR-associated transposases. **a,** Schematic of plasmid-to-plasmid transposition assay in human cells. **b,** Sanger sequencing confirmation of targeted transposition products after plasmids isolation from human cells and selected in *E. coli* (**panel a**), showing the expected insertion site position and presence of target-site duplication. **c,** Phylogenetic tree of Type I-F3 CRISPR-associated transposon systems adapted from previous work in the lab^39^, with labels of the homologs that were tested in human cells. **d,** Comparison of plasmid-to-plasmid integration efficiencies with eCAST-1 (*Vch*CAST) and. eCAST-2.1 (*Pse*CAST), as measured by qPCR. **e,** Optimization of eCAST-2 (*Pse*CAST) integration efficiencis by varying NLS placement and plasmid stoichiometries, etc., as described in **Supplementary Fig. 7**, yielded an approximate 6-fold increase in integration efficiencies. **f,** Amplicon sequencing reveals a strong preference for integration 49-bp downstream of the 3’ edge of the site targeted by the crRNA in T-RL integrants. **g,** Deletion experiments confirm the obligate requirement of each protein component, a targeting crRNA, and intact transposase active site (D220N mutation in TnsB, D458N mutation in TnsAB_f_) for successful integration. **h,** RNA-guided DNA integration functions with genetic payloads spanning 1–15 kb in size, transfected based on molar amount. **i,** RNA-guided DNA integration shows a strong sensitivity to mismatches across the entire 32-bp target site. Data were normalized to the perfectly matching (PM) crRNA, which exhibited an efficiency of 4.7 ± 1.8 %. Data in **d, e, g–i** are shown as mean ± s.d. for n = 3 biologically independent samples. Data in **d, e, g - i** are determined by qPCR.

After transfecting HEK293T cells with pDonor, pTarget, and all protein-RNA expression plasmids, purifying the plasmid mixture from cells, and using the mixture to transform *E. coli*, we observed the emergence of colonies that were chloramphenicol resistant, which outnumbered the corresponding colonies obtained from experiments using a non-targeting crRNA that did not match pTarget (**Supplementary Fig. 5a**). Encouraged by this result, we performed junction PCR on select colonies and obtained bands of the expected size, which subsequent Sanger sequencing confirmed were integration products arising from DNA transposition 49-bp downstream of the target site (**Fig. 4b**), as expected from our bacterial studies^38^. Further analyses of individual clones revealed the expected junction sequences across both the transposon left and right ends (**Supplementary Fig. 5b**). Next, we showed that the same products could be detected by nested PCR directly from HEK293T cell lysates (**Supplementary Fig. 5c**), and we developed a sensitive TaqMan probe-based qPCR strategy to quantify integration events from lysates by detecting site-specific, plasmid-transposon junctions (**Supplementary Fig. 5d**). Using this approach, we performed an initial optimization screen by varying the relative amounts of expression and pDonor plasmids and found that efficiencies were greatest with low levels of pTnsC and high levels of pTnsAB_f_ and pDonor (**Supplementary Fig. 5e**). Nevertheless, absolute efficiencies of plasmid-to-plasmid transposition with this engineered CAST system from *V. cholerae*, hereafter referred to as “engineered CAST-1” (eCAST-1), remained <0.1%, leading us to pursue other avenues for improved activity (**Supplementary Fig. 5e**).

We recently described the bioinformatic mining and experimental characterization of 18 new Type I-F CRISPR-associated transposons (denoted Tn*7000*–Tn*7017*), many of which exhibited high-efficiency and high-fidelity RNA-guided DNA integration in *E. coli* (**Fig. 4c**)^41^. We hypothesized that sampling from this diversity would uncover variants with improved activity in human cells, and thus embarked on a hierarchical screening approach to concentrate our efforts on the most promising systems (**Supplementary Fig. 6a**). Briefly, our scheme involved filtering based on robust activity in three key areas: (i) crRNA biogenesis by Cas6, assessed using our GFP knockdown assay; (ii) transposon DNA binding by TnsB, assessed using a tdTomato reporter assay; and (iii) transcriptional activation by TnsC-VP64, assessed using our mCherry reporter assay. In all cases, genes were human codon optimized, which was often necessary to achieve strong expression (**Supplementary Fig. 6b**), and tagged with NLS sequences on the same termini as for Tn*6677* (*Vch*CAST). We found that the majority of systems exhibited efficient crRNA biogenesis and transposon DNA binding activity that was similar to that observed with Tn*6677* (**Supplementary Fig. 6c,d**). Interestingly, of those systems selected for testing in transcriptional activation experiments, only Tn*7016* showed reproducible induction of mCherry expression, albeit at levels ∼8-fold lower than Tn*6677* (**Supplementary Fig. 6e**). We therefore decided to focus on Tn*7016* –– a 31-kb transposon from *Pseudoalteromonas* sp. S983 (*Pse*CAST) –– and next investigated its RNA-guided DNA integration activity.

After verifying that fusing TnsA and TnsB from *Pse*CAST with an internal NLS retained function, and optimizing the length of left and right transposon ends (**Supplementary Fig. 7a,b**), we repeated plasmid-to-plasmid transposition assays in HEK293T cells. Strikingly, the engineered *Pseudoalteromonas* CAST (eCAST-2.1) was ∼40-fold more active than eCAST-1 when tested under unoptimized conditions (**Fig. 4d**, **Supplementary Fig. 7c**). To further improve integration efficiencies, we systematically varied the design of the crRNA, location of NLS tags, and relative amounts of each expression plasmid; the resulting eCAST-2.2 yielded a further ∼6-fold improvement to reach levels of 3–5% integration, and PCR followed by Sanger or Illumina sequencing analysis confirmed the expected site of integration 49-bp downstream of the target (**Fig. 4e, f, Supplementary Fig. 7d–h**). Of note, these efficiencies were comparable to integration efficiencies achieved with BxbI under similar plasmid-to-plasmid conditions (**Supplementary Fig. 7i**). Peak integration occurred 4–6 days post-transfection, with the efficiency exhibiting sensitivity to both cell density and the choice of cationic lipid delivery method^62^ (**Supplementary Fig. 8a–c**). We also found that the observed integration efficiency was increased by >5-fold upon co-transfection of a GFP transfection marker and separately analyzing sorted cells exhibiting high GFP fluorescence levels, suggesting that activity was dependent not only the stoichiometry of the transfected plasmids but also the plasmid dosage across the population of cells (**Supplementary Fig. 8d,e**).

Next, we sought to confirm the genetic requirements for RNA-guided DNA integration and further investigate specificity. Transposition was strictly dependent on a targeting crRNA and the presence of all protein components, including an intact TnsB active site (**Fig. 4g**), and functioned with genetic payloads spanning 1–15 kb in size, albeit with a ∼3-fold decrease in efficiency with larger payloads (**Fig. 4h**). We generated a panel of mismatched crRNAs in which mutations were tiled along the length of the 32-nt guide, and found that activity was ablated regardless of the location (**Fig. 4i**), indicating a greater degree of discrimination than that observed in activation experiments utilizing *Vch*CAST in activation experiments or in *E. coli*^38^. We used an alternative qPCR approach to confirm that integration orientation for eCAST-2.2 was highly biased towards T-RL, as expected from prior bacterial integration data^41^ (**Supplementary Fig. 9a**). Finally, we utilized an NGS-based amplicon sequencing approach to quantify all integration events within a 10-bp window of the expected insertion site (**Supplementary Fig. 9b**) and performed droplet digital PCR (ddPCR) to further corroborate the quantitative data obtained from TaqMan qPCR (**Supplementary Fig. 9c**).

### RNA-guided DNA integration into the human genome

Following optimization efforts on episomal plasmid DNA integration with eCAST-2.2, we next turned our attention to reconstituting RNA-guided integration into endogenous genomic sites. We first screened a panel of guide sequences targeting the AAVS1 safe-harbor locus via a plasmid-to-plasmid integration assay, in which we cloned 32-bp target sites derived from AAVS1 into pTarget and leveraged existing assays to identify two active crRNAs that outperformed our original plasmid-specific crRNA (**Supplementary Fig. 10a**). When we tested the AAVS1 locus for genomic integration using a nested PCR strategy, we identified RNA-guided DNA integration products that again maintained the expected 49-bp distance dependence from the target site (**Fig. 5a**). However, detection was often not consistent across biological replicates, suggesting that integration efficiencies flirted with our limit of detection. We therefore applied an NGS-based amplicon sequencing method established in our prior plasmid-based assays, yielding reproducible efficiencies on the order of ∼0.005% (**Fig. 5b**).

**Fig. 5.**
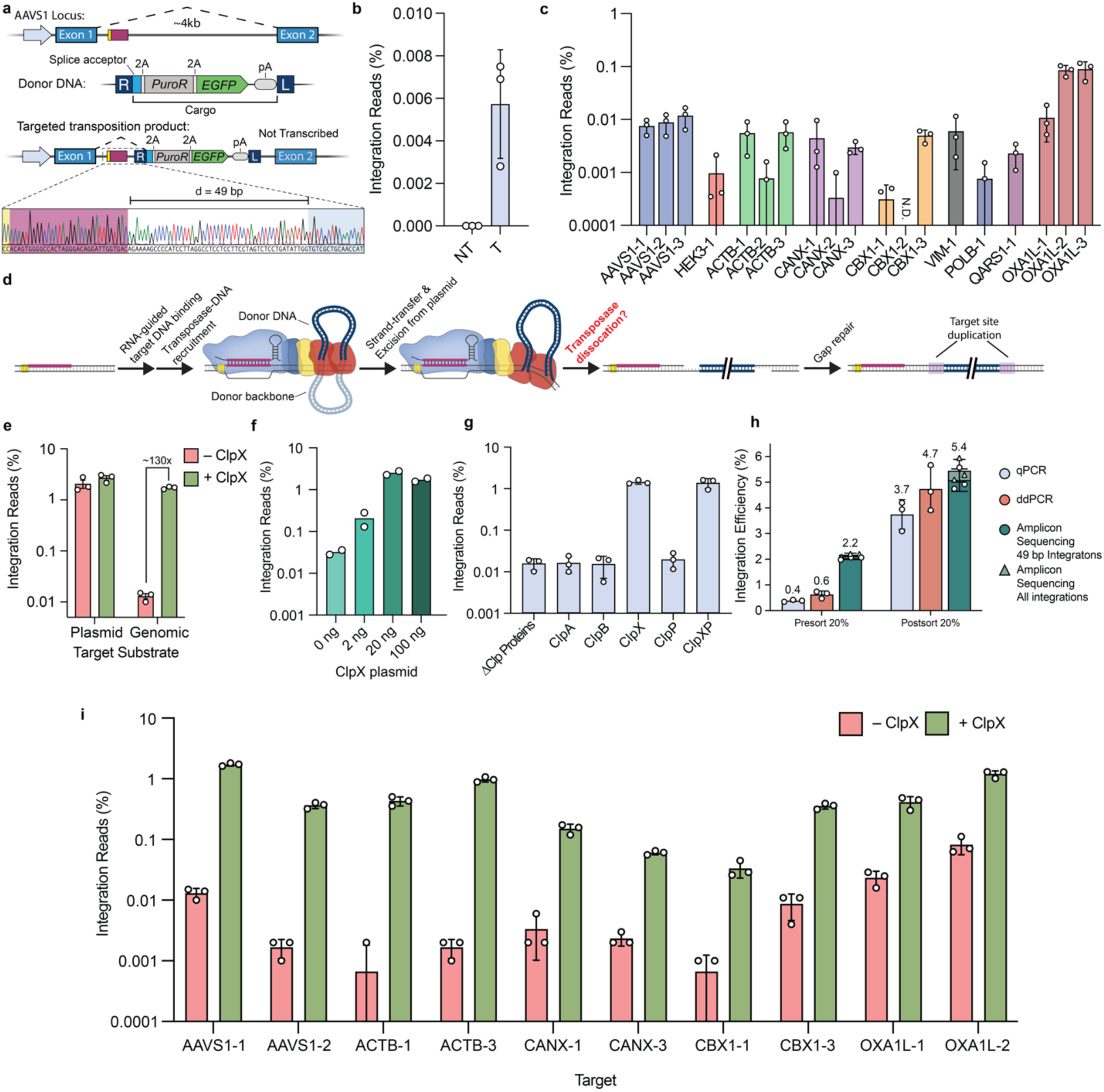
RNA-guided DNA integration into the human genome requires targeted degradation via ClpX. **a,** Sanger sequencing of nested PCR of genomic lysates in which eCAST-2.2 targeted the AAVS1 genome showed a junction product 49bp downstream of the target site targeted by crRNA12 (AAVS1-1), one of the optimal crRNAs screened in **Supplementary Fig. 10a. b,** Initial quantifications of genomic integration efficiencies at AAVS1-1. **c,** Integration efficiencies across multiple loci within human genome showed broadly limited efficiencies. Quantified integration efficiencies less than .0001% were not plotted, and “N.D.” represents a target site in which no integration events were detected across three biological replicates. **d,** Proposed steps required for successful targeted integration, including the downstream gap-repair needed for complete resolution of the integration product. **e,** Co-transfection of *Eco*ClpX specifically improves genomic, but not plasmid, integration efficiencies in human cells. **f,** Co-transfecting *Eco*ClpX at varied amounts directly impacts genomic integration efficiencies in human cells. **g,** Investigating the impact of various Clp proteins from *E. coli* on genomic integration efficiencies in human cells. **h,** Integration efficiencies for samples before and after FACS off of a fluorescent marker to select for transfected cells. The top 20% brightest cells were sorted and immediately harvested. Sorting increased in integration efficiencies, as measured by qPCR, ddPCR, and NGS. **i,** Integration efficiencies were investigated across multiple loci within the human genome with and without *Eco*ClpX. Quantified integration efficiencies less than .0001% were not plotted. Data in **b, c, e, g– i** are shown as mean ± s.d. for n = 3 biologically independent samples. Data in **f** are shown as mean for n = 2 biologically independent samples. Data in **b, c, e, f, g, and i** are quantified by amplicon sequencing.

We next targeted an additional 8 sites across the genome, with 1-3 crRNAs per locus, and detected integration at efficiencies that varied but were generally ∼0.01% (**Fig. 5c**). Attempts to increase the efficiency further through simplified delivery of a polycistronic QCascade expression vector, serial additions of extra NLS sequences, constitutive expression of the targeting machinery, inclusion of bacterial IHFa/b^63^, or phenotypic drug selection to enrich for integration events (**Supplementary Fig. 10b-f**) did not reduce the large, 100-1,000X discrepancy between observed integration efficiencies at plasmid versus genomic target sites. Although differences in chromatinization remained a distinct possibility, we hypothesized that the discrepancy might be due to potential toxicity of genomic integration intermediate products.

TnsB performs transesterification reactions to join the two ends of the transposon DNA to both strands of the target DNA with a 5-bp offset, leading to the generation of an initial product characterized by 5-nt gaps on either strand of the integrated DNA^64^. Subsequent gap repair involves gap fill-in by DNA polymerase and DNA ligase, resulting in the hallmark 5-bp target-site duplication (TSD), but these reactions require prior dissociation of the transpososome (**Fig. 5d**). We questioned whether incomplete dissociation of the post-transposition CAST complex might limit observed frequencies of genomic integration, perhaps leading to stalled replication forks or DNA repair pathways akin to crosslink-induced replication fork stalling^65–67^. Notably, these effects would likely be less deleterious on plasmid DNA substrates, since pTarget does not undergo active DNA replication and is not critical for cellular fitness. Our hypothesis was bolstered by prior studies demonstrating the extreme stability of the analogous post-transposition complex (PTC) in Tn*7* and Mu transposons^68–70^, and the requirement for an additional factor — ClpX — in active Mu PTC disassembly, gap repair, and phage propagation^68–72^. ClpX is a sequence-specific AAA+ ATPase that unfolds protein substrates by denaturing and translocating them through a central hexameric pore, and it recognizes degron tags that are often exposed only under certain conditions, allowing for sensitive regulation of protein unfolding and degradation^73^. In the case of MuA, ClpX recognizes a specific C-terminal motif, and the post-transposition complex undergoes a conformational rearrangement to expose additional MuA residues that enable more efficient ClpX binding, resulting in targeted unfolding and destabilization of the post-transposition complex^72, 74^. We hypothesized that CAST systems might also require bacterial ClpX, or some other accessory factor, for active mechanical disassembly of the PTC.

To test this, we co-transfected human cells with eCAST-2.2 components and a plasmid expressing NLS-tagged *E. coli* ClpX (*Eco*ClpX). Remarkably, genomic integration efficiencies increased by ∼100X in a dose-responsive manner when eCAST-2.2 was supplemented with *Eco*ClpX (referred to as eCAST-3 hereafter), whereas plasmid integration efficiencies were unaffected (**Fig. 5e,f**). To investigate if the effect was sequence-specific, we tested other bacterial disaggregases, including ClpA and ClpB, and found that ClpX was the only tested AAA+ ATPase factor that enhanced genomic integration (**Fig. 5g**). Moreover, ClpP, which functions as the peptidase component within the ClpXP protease complex, had no effect on integration, either alone or in combination with ClpX, suggesting that protein unfolding — but not protease degradation — is necessary for the enhancement effect (**Fig. 5g**). ClpX is highly conserved across bacterial species, and the homolog from *Pseudoalteromonas* (80% amino acid identity) also stimulated integration, albeit to a slightly lesser extent that *Eco*ClpX (**Supplementary Fig. 11a**); NLS-tagged human ClpX, which normally functions in the mitochondria, had no effect on integration (**Supplementary Fig. 11b**). Interestingly, genomic integration with eCAST-1 (*Vch*CAST) was reproducibly detectable in the presence of *Eco*ClpX or *Vch*ClpX but not in its absence, indicating a consistent effect across Type I-F CAST systems, though lower intrinsic activity of *Vch*CAST was observed similar to plasmid-to-plasmid integration assays (**Supplementary Fig. 11a**). Collectively, these results reveal PTC disassembly to be a critical bottleneck that limits integration into genomic target sites, and they identify ClpX as an accessory factor that acts directly to unfold one or more components within the CAST transpososome (**Supplementary Fig. 11c**). Future experiments will be needed to further dissect the mechanistic details.

Single-digit genomic integration efficiencies at the AAVS1 locus allowed us to explore other parameters of eCAST-3 design and delivery. We found that crRNAs functioned best with 33-nt spacers on both plasmid and genomic targets (**Supplementary Fig. 12a,b**), and that transfections could be simplified by placing the U6-driven crRNA cassette directly on pDonor without an adverse effect on activity (**Supplementary Fig. 12c**). Integration could be further improved with the appropriate selection of cationic lipid formulation (**Supplementary Fig. 12d**), or by selecting/sorting cells that were co-transfected with either a drug or fluorescent marker, with efficiencies reaching ∼5% as measured by amplicon-sequencing and ddPCR (**Fig. 5h, Supplementary Fig. 12e,f**). Importantly, we also carefully inspected our next-generation sequencing data to assess product purity at genomic sites of integration, looking specifically at whether unedited alleles showed any evidence of mutations, and whether edited alleles containing a transposon insertion harbored unexpected modifications. These analyses revealed an absence of indels above background (∼0.04% sequencing error) at unedited target sites, and an absence of detectable mutations surrounding genome-transposon junctions (**Supplementary Fig. 12g-i**), suggesting that CAST systems are less prone to the range of byproducts common to Cas9 nuclease and nickase-based approaches^23, 24^.

Lastly, we revisited previously targeted sites across the human genome and assessed integration efficiency to test the generalizability of ClpX enhancement (**Supplementary Fig. 13a**). Strikingly, we observed a 10–600-fold increase in integration efficiencies across all tested loci (**Fig. 5i**), with a consistent preference for insertions ∼49-bp downstream of the crRNA-matching target site (**Supplementary Fig. 13b**), as first reported in our *E. coli* studies^38, 39^.

## DISCUSSION

This work represents the first successful reconstitution of heteromeric, Tn7-like transposases in human cells, and establishes a strong foundation in the application of RNA-guided transposases for programmable, targeted DNA integration. Despite their molecular complexity, CRISPR-associated transposon (CAST) systems provide a direct route for inserting large genetic payloads at user-defined sites without generating DNA double-strand breaks, and without requiring multiple steps to combine editing and integrating modalities. While further advances will be necessary to make these systems broadly useful for research and therapeutic applications, our results progressing from eCAST-1 to eCAST-3 demonstrate feasibility of reconstituting multi-component editing pathways in human cells and highlight a robust pipeline to identify promising candidates for continued development.

We established functional assays to carefully assess each modular component of the *V. cholerae* Type I-F CAST system (eCAST-1), which revealed specific terminal tagging modifications that severely reduced, or in some cases, altogether eliminated activity (**Supplementary Fig. 2c**). Accordingly, the integration experiments in this study relied on transient delivery of multiple protein and RNA expression plasmids alongside the donor plasmid in a single co-transfection. Given the sensitivity of CAST systems to protein/complex stoichiometry, this approach reduces the fraction of cells that receive optimal distributions of each component. Moving forward, the increased amenability of eCAST-2 (i.e. *Pse*CAST) components to N- and C-terminal tagging, as compared to eCAST-1 (**Supplementary Fig. 7e**), together with structure-guided engineering and recent examples of naturally fused Class 1 complexes^75^, provide strong support for further streamlining the system into fewer molecular components while retaining its intrinsic properties. In addition, direct delivery of purified protein, RNA, and DNA components offers a particularly promising area of investigation, and electroporation of pre-assembled transpososomes comprising the transposon DNA and TnsAB_f_ may improve trafficking and/or co-localization of the donor genetic payload and transposase to the target site.

When we screened homologous systems, we observed a wide range of relative activities for crRNA maturation, transposon DNA binding, and TnsC-based transcriptional activation, indicating that each molecular step of the pathway may require independent optimization. For example, while components derived from Tn*6677* (*Vch*CAST) exhibited the strongest levels of activation, our assay for transposon DNA binding by TnsB revealed that homologs from Tn*7005* and Tn*7010* exhibited >200-fold activation in human cells (**Supplementary Fig. 6**). While we cannot exclude the possibility that the specific fusion constructs and reporter assay designs developed for this experiment fail to faithfully reflect TnsB activity, our results nevertheless suggest that none of the systems currently tested combine optimal activities in human cells across each molecular component.

In addition to the potential that RNA-guided transposases offer for DNA integration applications, we were excited to find that recruitment of the AAA+ ATPase TnsC, when fused with VP64 domains, stimulated robust levels of transcriptional activation at both plasmid and genomic target sites that were similar to levels achieved with dCas9-VPR fusion proteins. Recent structural and functional studies have demonstrated that TnsC homologs form ATP-dependent oligomers that assemble around dsDNA^58, 59, 76^, and we showed that TnsC is recruited to genomic loci in eukaryotic cells with high fidelity in a QCascade-dependent manner (**Fig. 3**), similarly to recent experiments performed in *E. coli*^49^. Thus, combining these molecular components, while foregoing the heteromeric transposase itself, reveals a potent strategy to assemble an intrinsically multimeric protein at user-defined target sites, for applications where multi-valency offers a considerable benefit. In addition to fusing TnsC to other activation or repression domains for control over gene expression levels, similar to existing CRISPRa and CRISPRi tools, we propose tethering epigenetic modifiers for DNA and/or histone modifications, or fluorescent proteins for higher signal-to-noise ratios for chromosomal loci imaging assays without requiring arrays of guide RNAs^77, 78^. Further, by also leveraging the multi-subunit nature of the QCascade complex, one could access more elaborate scaffolding approaches to recruit multiple functionalities to individual target sites simultaneously, such as by fusing effector domains to Cas8, Cas7, and/or TnsC in various combinatorial fashions.

Perhaps the most notable outcome of our study was the identification of bacterial ClpX as a novel accessory protein involved in CRISPR RNA-guided transposition (**Fig. 5, Supplementary Fig. 11**). The disparity we observed between integration efficiencies into episomal plasmid substrates versus genomic target sites inspired us to more carefully consider the importance of CAST transpososome disassembly to expose transposition product intermediates for gap fill-in and repair. Based on a careful review of the Tn7 and Mu transposon literature, we hypothesized that protein unfoldases might facilitate active mechanical dissociation of one or more CAST components, leading to the identification of ClpX as a stimulatory factor that increased integration efficiencies by more than two orders of magnitude (**Fig. 5**). Alongside another recent study that uncovered the surprising role of ribosomal protein S15 in transposition by Type V-K CAST systems^79^, our finding indicates that CAST systems may be more reliant on host proteins than previously appreciated, and that all chemical steps in the transposition pathway need to be critically evaluated. Although future efforts will focus on achieving transpososome destabilization and/or disassembly using strategies that obviate a need for additional factors, the combination of *Eco*ClpX and *Pse*CAST (eCAST-3) nonetheless allowed us to reach single-digit integration efficiencies across a range of genomic target sites (**Fig. 5i**), paving the way for further improvements.

CRISPR-based genome engineering tools have largely focused on single-protein effectors over the past decade, including Cas9, Cas12, and Cas13, because of the straightforward design of expression vectors, ease of viral delivery, and perceived simplicity in reconstitution. However, recent studies highlight the feasibility of transplanting more complex CRISPR-Cas effectors into eukaryotic cells, while retaining the ability to achieve high editing efficiencies and exploit novel enzymatic functionalities^42–45, 80^. Our work extends this paradigm further while leveraging a powerful new class of transposases that offer the promise of single-step insertion of large, multi-kilobase genetic payloads with the programmability afforded by RNA-guided CRISPR-Cas systems.

## METHODS

### Plasmid construction

Genes were human codon-optimized and synthesized by Genscript, and plasmids were generated using a combination of restriction digestion, ligation, Gibson assembly, and inverted (around-the-horn) PCR. All PCR fragments for cloning were generated using Q5 DNA Polymerase (NEB).

The CRISPR array sequence (repeat-spacer-repeat) for *Vch*CAST is as follows:

5’–GTGAACTGCCGAGTAGGTAGCTGATAAC–N_32_–GTGAACTGCCGAGTAGGTAGCTGATAAC–3’ where N_32_ represents the 32-nt guide region. The sequence of the mature crRNA is as follows: 5’–CUGAUAAC–N_32_–GUGAACUGCCGAGUAGGUAG–3’

The CRISPR array sequence (repeat-spacer-repeat) for *Pse*CAST is as follows:

5’–GTGACCTGCCGTATAGGCAGCTGAAAAT–N_32_–GTGACCTGCCGTATAGGCAGCTGAAAAT–3’ where N_32_ represents the 32-nt guide region. The sequence of the mature crRNA is as follows: 5’–CUGAAAAU–N_32_–GUGACCUGCCGUAUAGGCAG–3’

We also used ‘atypical’ repeats^41, 81^ for *Pse*CAST (unless otherwise mentioned) to reduce the likelihood of recombination during cloning. For these variant CRISPR arrays, the repeat-spacer-repeat sequence is as follows:

5’–GTGACCTGCCGTATAGGCAGCTGAAGAT–N_32_–TAATTCTGCCGAAAAGGCAGTGAGTAGT–3’ where N_32_ represents the 32-nt guide region. The sequence of the mature crRNA is as follows: 5’–CUGAAGAU–N_32_–UAAUUCUGCCGAAAAGGCAG–3’

Where noted, we modified the 32-nt guide region to have varying lengths. The repeat sequences flanking the guide region were not modified in these experiments.

Clp proteins from the *E. coli* genome were PCR amplified from BL21 DE3 cells with primers that specifically amplified the open reading frame of the indicated protein and cloned into pcDNA3.1 expression vectors with an N-terminal bipartite-NLS tag. ClpX sequences from *E. coli*, *Pseudoalteromonas sp.*, and *V. cholerae* were then codon-optimized by Genscript and ordered as Twist fragments to be cloned into pcDNA3.1 expression vectors with an N-terminal bipartite-NLS tag.

### *E. coli* culturing and general transposition assays

Chemically competent *E. coli* BL21(DE3) cells carrying pDonor, pDonor and pTnsABC, or pDonor and pQCascade, were prepared and transformed with 150–250 ng of pEffector, pQCascade, or pTnsABC, respectively. Transformations were plated on agar plates with the appropriate antibiotics (100 µg/ml spectinomycin, 100 µg/ml carbenicillin, 50 µg/ml kanamycin) and 0.1 mM IPTG. For bacterial transposition assays investigating *Pse*CAST activity, cells were co-transformed with pEffector and pDonor. Cells were incubated for 18–20 h at 37 ℃ and typically grew as densely spaced colonies, before being scraped, resuspended in LB medium, and prepared for subsequent analysis. A full list of all plasmids used for transposition experiments is provided in **Supplementary Table 1**, and a list of crRNAs used is provided in **Supplementary Table 2**.

### *E. coli* qPCR analysis of transposition products

The optical density of resuspended colonies from our transposition assays was measured at 600 nm, and approximately 3.2 × 10^8^ cells (the equivalent of 200 µl of OD600 = 2.0) were pelleted by centrifugation at 4,000 x g for 5 min. The cell pellets were resuspended in 80 µl of H_2_O, before being lysed by incubating at 95 °C for 10 min in a thermal cycler. The cell debris was pelleted by centrifugation at 4,000 x g for 5 min, and 5 µl of lysate supernatant was removed and serially diluted in water to generate 20- and 500-fold lysate dilutions for qPCR analysis.

Integration in the T-RL orientation was measured by qPCR by comparing Cq values of a tRL-specific primer pair (one transposon- and one genome-specific primer) to a genome-specific primer pair that amplifies an *E. coli* reference gene (*rssA*). Transposition efficiency was then calculated as 2^ΔCq^, in which ΔCq is the Cq difference between the experimental reaction and the reference reaction. A full list of oligos used for integration efficiency measurements is provided in **Supplementary Table 3**. qPCR reactions (10 µl) contained 5 µl of SsoAdvanced Universal SYBR Green Supermix (BioRad), 1 µl H2O, 2 µl of 2.5 µM primers, and 2 µl of 500-fold diluted cell lysate. Reactions were prepared in 384-well clear/white PCR plates (BioRad), and measurements were performed on a CFX384 Real-Time PCR Detection System (BioRad) using the following thermal cycling parameters: polymerase activation and DNA denaturation (98 °C for 3 min), and 35 cycles of amplification (98 °C for 10 s, 59 °C for 1 min).

### Mammalian cell culture and transfections

HEK293T cells were cultured at 37 °C and 5% CO_2_. Cells were maintained in DMEM media with 10% FBS and 100 U/mL of penicillin and streptomycin (Fisher Scientific). The cell line was authenticated by the supplier and tested negative for mycoplasma.

Cells were typically seeded at approximately 100,000 cells per well in a 24-well plate (Eppendorf or Fisher Scientific) coated with poly-D-lysine (Fisher Scientific), 24 hours prior to transfection. Cells were transfected with DNA mixtures and 2 µl of Lipofectamine 2000 (Fisher Scientific), per the manufacturer’s instructions. Transfection reactions typically contained between 1µg and 1.5µg of total DNA. For detailed transfection parameters specific to distinct assays, please refer to the sections below.

### Western immunoblotting and nuclear/cytoplasmic fractionation

Cells were transfected with epitope-tagged protein expression plasmids. Approximately 72 hours after transfection, cells were washed with PBS and harvested using Cell Lysis Buffer (150 mM NaCl, 0.1% Triton X-100, 50mM Tris-HCl pH 8.0, Protease inhibitor (Sigma Aldrich)). For nuclear and cytoplasmic fractionation experiments, cells were harvested using Cell Lysis Buffer (Thermo Fisher Scientific) per the manufacturer’s instructions. Proteins were separated by SDS-PAGE and transferred to a PVDF membrane (Fisher Scientific). The membrane was then washed with TBS-T (50mM Tris-Cl, pH 7.5, 150mM NaCl, .1% Tween-20) and blocked with blocking buffer (TBS-T with 5% w/v BSA). Membranes were then incubated with primary antibodies overnight at 4°C in blocking buffer. Membranes were then washed and incubated with secondary antibodies at room temperature for one hour. Membranes were again washed and then developed with SuperSignal West Dura (Thermo Fisher). Antibodies used for immunostaining are listed in **Supplementary Table 4**.

### HEK293T fluorescent reporter assays and flow cytometry analysis and sorting

HEK293T cells were seeded at approximately 50,000 cells per well in a 24-well plate coated with poly-D-lysine 24 hours prior to transfection. For Cas6-mediated RNA processing assays, cells were co-transfected with 300 ng of GFP-reporter plasmid, 300 ng of Cas6 expression plasmid, and 10 ng of an mCherry expression plasmid (as a transfection marker). In negative control experiments, cells were transfected with 300 ng of a dCas9 expression plasmid instead of a Cas6 expression plasmid to control for possible expression burden or squelching. For transcriptional activation assays, cells were co-transfected with 60 ng of reporter plasmid, 20 ng of a plasmid encoding an orthogonal fluorescent protein (as a transfection marker), and the additional indicated plasmids. In separate wells, cells were transfected with 100 ng of Cas9-based transcriptional activators and 50 ng of either a non-targeting or targeting sgRNA as positive controls^82^.

DNA mixtures were transfected using 2 µl of Lipofectamine 2000 (Fisher Scientific), per the manufacturer’s instructions. Approximately 72–96 hours after transfection, cells were collected for assay by flow cytometry. Transfected cells were analyzed by gating based on fluorescent intensity of the transfection marker relative to a negative control, as previously described^82^. For assays that involved cell sorting, cells were transfected with a GFP expression plasmid and collected 4 days after transfection. A BD FACS Aria flow cytometer was used to sort cells and obtain flow cytometry data. Cells with the top 20% brightest GFP fluorescence were sorted by 5% increments into 4 bins. Cells were immediately harvested after sorting, as detailed below.

### HEK293T genomic activation and RT-qPCR analysis

HEK293T cells were seeded at approximately 50,000 cells per well in a 24-well plate coated with poly-D-lysine 24 hours prior to transfection. Cells were co-transfected as described above, with the following *Vch*CAST components: 100 ng pTnsAB_f_ , 50 ng pTnsC-VP64, 50 ng pTniQ, 50 ng pCas6, 250 ng pCas7, 50 ng pCas8, and 62.5 ng each of 4 targeting crRNAs for *TTN*, *MIAT*, and *ASCL1* (or 83.3 ng each of 3 targeting crRNAs for *ACTC1*) (pCRISPR). In control experiments, cells were co-transfected with 100 ng of either pdCas9-VP64 or pdCas9-VPR plasmid, 62.5 ng each of 4 targeting sgRNAs for *TTN* (psgRNA), and a pUC19 plasmid to standardize transfected DNA amounts; see **Supplementary Table 2** for crRNAs and sgRNAs used. Cells were harvested 72 hours after transfection using the RNeasy Plus Mini Kit (Qiagen), according to the manufacturer’s instructions. cDNA was subsequently synthesized using the iScript cDNA Synthesis Kit (BioRad) using 1000 ng of RNA in a 20 uL reaction. Gene-specific qPCR primers^61^ were designed to amplify an approximately 180-250 bp fragment to quantify the RNA expression of each gene, and a separate pair of primers was designed to amplify *ACTB* (beta-actin) reference gene for normalization purposes. A comprehensive list of oligonucleotides used in the study is available in **Supplementary Table 3**.

qPCR reactions (10 µl) contained 5 µl of SsoAdvanced Universal SYBR Green Supermix (BioRad), 2 µl H_2_O, 1 µl of 5 µM primer pair, and 2 µl of cDNA diluted 1:4 in H_2_O. Reactions were prepared in 384-well white PCR plates (BioRad), and measurements were performed on a CFX384 Real-Time PCR Detection System (BioRad) using the following thermal cycling parameters: polymerase activation and DNA denaturation (98 °C for 2 min), 40 cycles of amplification (95 °C for 10 s, 60 °C for 30 s), and terminal melt-curve analysis (65–95 °C in 0.5 °C per 5 s increments). Each condition was analyzed using three biological replicates, and two technical replicates were run per sample. Normalized gene activation was calculated as the ratio of the 2^-Δ*C*q^ of the targeting samples to the non-targeting samples, in which Δ*C*q is the *C*q difference between the experimental gene primer pair and the reference gene primer pair.

### Chromatin Immunoprecipitation

For ChIP-seq analysis experiments, HEK293T cells were seeded at approximately 1,500,000 cells per well in a 10 cm dish coated with poly-D-lysine 24 hours prior to transfection. Cells were co-transfected as described above with the following eCAST-1 components: 1.5 ug p3xFLAG-TnsC, 1.5 ug pTniQ, 1.5 ug pCas6, 7.5 ug pCas7, 1.5 ug pCas8, and 3 ug of either a targeting (*TTN* crRNA 1) or non-targeting crRNA. See **Supplementary Table 2** for crRNAs used. 72 hours after transfection, cells were washed with DPBS with no calcium or magnesium (Fisher Scientific), harvested using TrypLE (Fisher Scientific), and neutralized with culture media. The resuspended cells were pelleted by centrifugation at 300 x g for 5 minutes, and the supernatant was aspirated. The pellets were processed as described previously^49, 83, 84^. In brief, pellets were resuspended in 1% freshly made formaldehyde (Thermo Fisher Scientific in DPBS and shaken gently for 10 minutes. Fixation was quenched by adding 2.5 M glycine, for a final concentration of 125 mM glycine, and shaking cells gently for 5 minutes. Cells were pelleted as described above, washed with cold DPBS, pelleted, resuspended in DPBS and 1x cOmplete EDTA free protease inhibitors (Sigma Aldrich), pelleted, flash frozen in liquid nitrogen, and stored at −80 °C.

On the day of sonication, the cross-linked pellets were resuspended in 1 mL of Lysis Buffer 1 (50 mM HEPES-KOH, 140 mM NaCl, 1 mM EDTA, 10% glycerol, 0.5% NP-40, 0.25% Triton X-100) and 1X protease inhibitors and rotated for 10 minutes. Cells were pelleted at 1350 g for 5 minutes. Pellets were resuspended in 1 mL of Lysis Buffer 2 (10 mM Tris-HCl, 200 mM NaCl, 1 mM EDTA, 0.5 mM EGTA) and 1X protease inhibitors and rotated for 10 minutes before being pelleted at 1350 g for 5 minutes. Pellets were resuspended in 900 uL of Lysis Buffer 3 (10 mM Tris-HCl, 100 mM NaCl, 1 mM EDTA, 0.5 mM EGTA, 0.1% Na-Deoxycholate, 0.5% N-lauroylsarcosine), 100 uL of 10% Triton X-100, and 1X protease inhibitors. All steps took place at 4 °C.

The resuspended cells were transferred to 1 ml milliTUBE AFA Fiber (Covaris) and sonicated on M220 Focused-ultrasonicator (Covaris) under the following SonoLab 7.2 settings: minimum temperature 4°C, set point 6 °C, maximum temperature 7 °C, Peak Power 75.0, Duty Factor 10.0, Cycles/Burst 200, sonication time 490 seconds. Sonicated cell lysate was centrifuged at 20,000 g for 10 minutes at 4 °C. The supernatant was transferred to a new tube, and 5% was saved as the input sample. The remaining supernatant was incubated with Dynabeads Protein G (Thermo Fisher Scientific) that were bound to the monoclonal anti-Flag M2 antibody (Sigma-Aldrich) the day before sonication by overnight rotating at 4 °C, and the lysate-Dynabead mixture was rotated overnight at 4 °C.

The samples were washed three times each with low salt buffer (150 mM NaCl, 0.1% SDS, 1% Triton X-100, 1 mM EDTA, 50 mM Tris HCl), high salt buffer (550 mM NaCl, 0.1% SDS, 1% Triton X-100, 1 mM EDTA, 50 mM Tris HCl), and LiCl buffer (150 mM LiCl, 0.5% Na-deoxycholate, 0.1% SDS, 1% Nonidet P-40, 1 mM EDTA, 50 mM Tris HCl) on a magnetic stand at 4 °C. The samples were washed with 1 mL of TE buffer (1 mM EDTA, 10 mM Tris HCl) with 50 mM NaCl and centrifuged at 960 g for 3 minutes at 4 °C. The supernatant was carefully aspirated with a pipette. 210 uL of elution buffer (1% SDS, 50 mM Tris HCl, 10 mM EDTA, 200 mM NaCl) was added to samples and incubated for 30 minutes at 65 °C. Samples were centrifuged for 1 minute at 16,000 g at room temperature, and 200 uL of supernatant was incubated overnight at 65 °C. The input sample was diluted in 150 uL of elution buffer and also incubated overnight at 65 °C. 0.5 uL of 10 mg/mL RNase was added, and samples were incubated for 1 hour at 37 °C. 2 uL of 20 mg/mL Proteinase K were added, and samples were incubated for 1 hour at 55 °C. The DNA was recovered by the QiaQUICK PCR Purification Kit (Qiagen) and DNA was eluted in 50 uL of water for downstream analysis.

### ChIP-seq Sample Preparation

Sample DNA concentration was determined by the DeNovix dsDNA High Sensitivity Kit. Illumina libraries were generated using the NEBNext Ultra II Dna Library Prep Kit for Illumina (NEB), as described previously^49^. Sample concentrations were normalized such that 12 ng of DNA in each condition was used for library preparation. The concentration of DNA was determined for pooling using the DeNovix dsDNA High Sensitivity Kit. Illumina libraries were sequenced in paired-end mode on the Illumina NextSeq platforms with automated demultiplexing and adaptor trimming. For each ChIP-seq sample, 75-bp paired end reads were obtained and between 9.5 and 18.9 million uniquely mapped fragments were analyzed.

### ChIP-seq analysis

ChIP-seq data were processed using CoBRA v2.0^85^ with modifications as follows. Each experimental condition (TnsC with *TTN-*targeting gRNA or TnsC with non-targeting [NT] gRNA) was processed with three biological replicate ChIP samples and one corresponding non-immunoprecipitated input sample. Reads were aligned to the hg38 human reference genome using BWA-MEM with default settings. Reads were sorted and indexed using SAMtools^86^, and multi-mapping reads with a MAPQ score < 1 were removed using the samtools view command. Peaks were called using MACS2 v2.2.6^87^. The callpeak function was executed in paired-end mode with the following parameters: -g 2.7e9 -q 0.0001 --keep-dup auto --nomodel. Input samples were used as controls for peak calling. Bedgraph files for each sample with pileup information in signal per million reads (SPMR) were generated with the --SPMR and -B subcommands of MACS2 callpeak and were converted to bigwig files using bedGraphToBigWig. ChIP-seq signal at individual genomic loci was visualized with IGV^88^. Reads mapping to the Y chromosome or the mitochondrial genome were removed prior to downstream analysis.

A consensus list of peaks for each experimental condition was identified using bedtools v2.30.0^89^. First, peak files for the three replicates were concatenated and sorted and overlapping peaks were merged. Then, peaks appearing in fewer than three replicates were removed. Blacklisted regions of the genome defined by the ENCODE Consortium were also removed^90^. The consensus lists for the conditions were then intersected to identify peaks exclusive to either condition (bedtools intersect -v) or peaks shared by both conditions (bedtools intersect -u). Differential binding analysis was performed using DiffBind v3.6.5^91^ to compare ChIP-seq read density between the two conditions in the regions defined by their consensus peak lists. Reads were counted using dba.count with the following arguments: summits = F, bUseSummarizeOverlaps = T, bRemoveDuplicates = F, bSubControl = F. Read counts were normalized to account for differences in sequencing depth between samples. Normalized read counts were passed to DESeq2 to calculate the mean across conditions, as well as fold change and q-value (using the Benjamini-Hochberg procedure) between conditions, for each peak. The result of differential binding analysis was visualized using ggplot2.

Heatmaps of ChIP-seq signal intensity over peaks exclusive to the *TTN* gRNA condition were plotted using deepTools v3.3.2^92^. Score matrices were generated using computeMatrix in reference-point mode. Peaks were sorted in descending order by mean signal over 2 kb windows around peak centers before plotting using plotHeatmap.

For manual inspection of potential off-target sites, a custom script was used to identify genomic loci with high similarity to the *TTN* spacer sequence. Other than the *TTN* locus itself, no loci with fewer than 5 mismatches were identified. TnsC ChIP-seq signal at the 5 most similar loci was visualized with IGV.

### HEK293T integration assays

For assays in which plasmids were isolated and used to transform bacteria, HEK293T cells were transfected with requisite eCAST-1 expression plasmids, a pDonor that contained a non-replicative origin of replication (R6K), a pTarget plasmid, and a crRNA expression plasmid (pCRISPR) that either encoded a non-targeting crRNA or a crRNA targeting pTarget. 72 hours after transfection, cells were thoroughly washed with PBS, harvested using TrypLE (Fisher Scientific), neutralized with culture media, and pelleted. After removal of supernatant, transfected plasmids were harvested using Qiagen Miniprep columns per the manufacturer’s instructions, and further concentrated using the Qiagen MinElute column. Of this final purified plasmid mixture, 1 µl was used to electroporate NEB 10-beta electrocompetent E. coli cells (NEB) per the manufacturer’s instructions. After recovery at 37 °C, cells were plated onto LB-agar plates containing chloramphenicol. Chloramphenicol-resistant colonies were then replated onto LB-agar plates containing both chloramphenicol and kanamycin, and doubly-resistant colonies were harvested for genotypic analyses.

For all other integration assays, HEK293T cells were counted using a Countess 3 Cell Counter and seeded at 20,000 cells per well, unless otherwise specified, in a 24-well plate coated with poly-D-lysine 24 hours prior to transfection. Cells were transfected using plasmid DNA mixtures and 2 µl of Lipofectamine 2000, per the manufacturer’s instructions. For eCAST-1 transposition assays, HEK293T cells were transfected with the following optimized *Vch*CAST components, unless otherwise stated: 300 ng of pTnsAB_f_, 25 ng of pTnsC, 100ng each of pTniQ, pCas6, pCas7, pCas8, 200 ng of pDonor, 100 ng pTarget, and 100 ng of a targeting or non-targeting crRNA (pCRISPR). For eCAST-2 transposition assays, HEK293T cells were transfected with the following *Pse*CAST components, unless otherwise specified: 200 ng of pTnsAB_f_, 50 ng each of pTnsC, pTniQ, pCas6, pCas7, and pCas8, 200 ng of pDonor, and 100 ng of pTarget and a targeting or non-targeting crRNA (pCRISPR). When a QCascade polycistronic expression vector was used (pQCas), 75ng was transfected. For eCAST-3 transposition assays, eCAST-2 conditions were used with pQCas, and 20ng of pClpX was co-transfected as well (unless otherwise noted). All eCAST-3 transposition assays utilized puromycin selection (unless otherwise noted, see below for puromycin conditions), as constitutive ClpX expression led to visible toxicity independent of CAST machineries. A full list of plasmids and crRNAs used is available in **Supplementary Table 1** and **Supplementary Table 2**, respectively. Unless otherwise stated, cells were cultured for 4 days after transfection. Cells were washed with DPBS with no calcium or magnesium (Fisher Scientific), harvested using TrypLE (Fisher Scientific), and neutralized with culture media. 20% of the resuspended cells were pelleted by centrifugation at 300 x g for 5 minutes, and the supernatant was aspirated. Cell pellets were resuspended in 50 µL of Quick Extract (Lucigen), and genomic DNA was prepared per the manufacturer’s instructions.

For assays that utilized puromycin selection, HEK293T cells were transfected as described above with *Pse*CAST component plasmids and an additional 20 ng of puromycin resistance expression plasmid (as a transfection marker). Media was changed 24 hours after transfection, and selection with 1 µg/mL of puromycin was started on half of the samples. Cells were harvested using Quick Extract (Lucigen) per the manufacturer’s instructions beginning at 2 days after transfection until 6 days after transfection, with or without puromycin selection. For plasmid-based assays that utilized cell sorting, HEK293T cells were transfected as described above with *Pse*CAST component plasmids and an additional 5 ng of GFP expression plasmid (as a transfection marker). 4 days after transfection, the GFP positive cells with the brightest mean fluorescence intensity were sorted in 4 bins of 5% increments to encompass the 20% brightest cells, and were immediately harvested as described above. For genomic assays that utilized cell sorting, HEK293T cells were seeded at approximately 100,000 cells in 6 well plates coated with poly-D lysine 24 hours before transfection. Cells were transfected with the following eCAST-3 components: 1000 ng each of pTnsAB_f_ and pDonor, 250 ng of pTnsC, 375 ng of polycistronic pCas7-Cas8-Cas6- TniQ, 20 ng of pGFP, 100 ng of pClpX, and 500 ng of a targeting crRNA (pCRISPR). 4 days after transfection, the top 20% of GFP positive cells with the brightest mean fluorescence intensity were sorted and immediately harvested, as described above. For genomic integration assays, cells were harvested by previously described assays, using 100μl of freshly prepared lysis buffer (10 mM Tris-HCl, pH 7.5; 0.05% SDS; 25 μg/ml proteinase K (ThermoFisher Scientific) directly into each well of the tissue culture plate. The genomic DNA mixture was incubated at 37 °C for 1–2 h, followed by an 80 °C enzyme inactivation step for 30 min^22^.

For assays that utilized cargo sizes ranging from 798 bp to 15 kb, HEK293T cells were transfected as described above with *Pse*CAST component plasmids, except the 5 kb, 10 kb, and 15 kb pDonor plasmids were transfected in molar equivalents to the 798 bp pDonor (∼406 fmol), to account for the size difference between donor plasmids. For assays that utilized amplicon deep sequencing, HEK293T cells were transfected as described above, with a pDonor plasmid that contained a primer binding site immediately downstream of the right transposon end that matched a primer binding site present in the unedited pTarget plasmid. Cells were harvested 4 days after transfection.

### Nested PCR analysis of transposition assays

DNA amplification was performed by PCR using Q5 Hot Start High-Fidelity DNA Polymerase (NEB) following the manufacturer’s protocol. In brief, 1 µL of cell lysate was added to a 25 µL PCR reaction. Thermocycling conditions were as follows: 98 °C for 45 seconds, 98 °C for 15 seconds, 66 °C for 15 seconds, 72 °C for 10 seconds, 72 °C for 2 minutes, with steps 2–4 repeated 24 times. The annealing temperature was adjusted depending on primers used. 1 µL of the first PCR reaction served as the template for a second 25 µL PCR reaction that was run under the same thermocycling conditions. Primer pairs contained one target-specific primer and one transposon-specific primer, and the primers used in the second PCR reaction generated a smaller amplicon than the first reaction (see **Supplementary Table 3** for oligonucleotides used in this study). PCR amplicons were resolved by 1–2% agarose gel electrophoresis and visualized by staining with SYBR Safe (Thermo Scientific). Negative control samples were always analyzed in parallel with experimental samples to identify mis-priming products, some of which presumably result from the analysis being performed on crude cell lysates that still contain the pDonor and target-site DNA.

### qPCR analysis of plasmid-to-plasmid and genomic transposition products

Transposition-specific qPCR primers were designed to amplify a ∼140-bp fragment to quantify transposition efficiency. Primer pairs were designed to span a transposition junction, with the forward primer annealing to pTarget, or the genome, and the reverse primer annealing within the transposon. Additionally, a custom 5’ FAM-labeled, ZEN/3’ IBFQ probe (IDT) was designed to anneal to the target-transposon junction. A separate pair of primers and a SUN-labeled, ZEN/3’ IBFQ probe (IDT) were designed to amplify a distinct segment of the target plasmid, or the human genome, for efficiency calculation purposes. For a full list of oligonucleotides used in qPCR, please refer to **Supplementary Table 3**.

Probe-based qPCR reactions (10 uL) contained 5 uL of TaqMan Fast Advanced Master Mix, 0.5 uL of each 18 uM primer pair, 0.5 uL of each 5 uM probe, 1 uL of H_2_O, and 2 uL of ten-fold diluted cell lysate for plasmid-based transposition samples, or 2 uL of five-fold diluted cell lysate for genomic transposition samples. Reactions were prepared in 384-well white PCR plates (BioRad), and measurements were performed on a CFX384 Real-Time PCR Detection System (BioRad) using the following thermal cycling parameters: polymerase activation (95 °C for 10 minutes) and 50 cycles of amplification (95 °C for 15 seconds, 59.5 °C for 1 minute). Each condition was analyzed using either two or three biological replicates, and two technical replicates were run per sample. Baseline threshold ratios were manually adjusted to be 1:1 for the reference primer pair to the transposition primer pair. Transposition efficiency was calculated as a percentage as 2^-Δ*C*q^ times 100, in which Δ*C*q is the Cq difference between the reference primer pair and the transposition primer pair.

To analyze the frequency of left-right insertion (tLR) versus right-left insertion (tRL) of the *Pse*CAST transposon in plasmid-based assays, transposition-specific qPCR primers were designed to span the tLR transposition junction, in addition to the primer pairs used for tRL integration and the reference amplicon in the probe-based qPCR analysis described above. qPCR reactions (10 uL) contained 5 µl of SsoAdvanced Universal SYBR Green Supermix (BioRad), 2 µl H_2_O, 1 µl of 5 µM primer pair, and 2 µl of ten-fold diluted cell lysate. Reactions were prepared in 384-well white PCR plates (BioRad), and measurements were performed on a CFX384 Real-Time PCR Detection System (BioRad) using the following thermal cycling parameters: polymerase activation and DNA denaturation (98 °C for 2 min), 50 cycles of amplification (95 °C for 10 s, 59.5 °C for 20 s), and terminal melt-curve analysis (65–95 °C in 0.5 °C per 5 s increments). Each condition was analyzed using three biological replicates, and two technical replicates were run per sample.

### ddPCR analysis of transposition products

During harvesting of HEK293T plasmid-based transposition assays, 50% of the resuspended cells were reserved during lysate generation. 500 µL of resuspended cells were pelleted by centrifugation at 300 x g for 5 minutes. The supernatant was aspirated, and DNA was extracted from cell pellets using the Qiagen DNeasy Blood and Tissue Kit (Qiagen). DNA was eluted in H_2_O and diluted to a concentration of 2.5 ng/µL. For genomic transposition assays, crude cell lysate, generated as described above, was purified using two-sided AMPure XP beads (Beckman Coulter) as follows^49^: 45 uL of AMPure XP beads were added to 20-80 uL of genomic lysate and incubated for 5 minutes before being placed on a magnetic PCR rack for 5 minutes. The supernatant was aspirated, and the beads were washed twice with 80% ethanol. The beads were dried for 5 minutes, then 25 uL of water was added to resuspend the beads. The suspension was incubated for 10 minutes off the magnetic rack, then placed back on the rack for 5 minutes. The supernatant was transferred to a new tube.

ddPCR was performed with the same primers and probes as detailed above for plasmid-to-plasmid transposition analysis and genomic transposition assays with the exception of OXA1L-2 target site, which was not quantified via qPCR. For a full list of oligonucleotides used in ddPCR, please refer to **Supplementary Table 3**. Plasmid-based ddPCR reactions (20 µL) contained 10 µL of ddPCR Supermix for Probes (Biorad), 1 µL of each 5 µM probe, 1 µL of each 18 µM primer pair, 5 units of HindIII (NEB), 4.13 µL of H_2_O, and 2 µL of 2.5 ng/µL DNA. Genomic ddPCR reactions (20 µL) contained 10 µL of ddPCR Supermix for Probes (Biorad), 1 µL of each 5 µM probe, 1 µL of each 18 µM primer pair, 5 units of HindIII (NEB), and 6.33 uL of purified DNA, ranging from ∼6 ng to ∼500 ng. Reactions were assembled at room temperature, and droplets were generated using the Biorad QX200 Droplet Generator according to the manufacturer’s instructions. Thermocycling was performed on a Biorad C1000 Touch Thermocycler with the following parameters: enzyme activation (95 °C for 10 minutes), 40 cycles of amplification (94 °C for 30 second, 61.5 °C for 1 minute) and enzyme deactivation (98 °C for 10 minutes). After thermocycling, droplets were hardened at 4 °C for 2 hours. Droplets were analyzed using the QX200 Droplet Reader according to the manufacturer instructions. Transposition percentages were calculated as the number of FAM positive molecules divided by the number of SUN/VIC positive molecules times 100.

### Preparation of amplicons for NGS analysis

PCR-1 products were generated as described above, except primers contained universal Illumina adaptors as 5′ overhangs (**Supplementary Table 3**) and the cycle number was reduced to 15 for plasmid-to-plasmid integration assays, and 25 for genomic integration assays. Additionally, up to 5 degenerate nucleotides was placed between the primer binding site and the Illumina adaptor 5’ overhang to improve library diversity when sequencing. 1µl of lysate per 10µl of overall PCR reaction was used; plasmid-to-plasmid integration assays were 20µl PCR reactions, while genomic integration assays were 250µl PCR reactions to sample sufficient alleles. These products were then diluted 20-fold into a fresh polymerase chain reaction (PCR-2) containing indexed p5/p7 primers and subjected to 10 additional thermal cycles using an annealing temperature of 65 °C. After verifying amplification by analytical gel electrophoresis, barcoded reactions were pooled and resolved by 2% agarose gel electrophoresis, DNA was isolated by Gel Extraction Kit (Qiagen), and NGS libraries were quantified by qPCR using the NEBNext Library Quant Kit (NEB). Illumina sequencing was performed using the NextSeq platform with automated demultiplexing and adaptor trimming (Illumina).

To determine the integration site efficiencies and distributions, a 10-bp sequence immediately downstream of the forward primer is identified to verify that an amplicon is an on-target amplicon. If a 10-bp transposon-end sequence is present in the amplicon, it is counted as an integration event and the distance is calculated from the start of the transposon to the PAM-distal end of the target sequence. % of reads that contained integration events are calculated as the proportion of integration reads to total on-target amplicons. Histograms were plotted after compiling these distances across all the reads within a given library.

### Data availability

Sequencing data has been deposited in the National Center for Biotechnology Information Sequence Read Archive under GEO accession GSE223174.

## ACKNOWLEDGMENTS

We thank N. Jaber and S.R. Pesari for laboratory support, L.F. Landweber for qPCR instrument access, the Columbia University Institute for Genomic Medicine for ddPCR instrument access, W. Frankel for thermocycler instrument access, the Columbia Stem Cell Initiative Flow Cytometry Core and M. Kissner and R. Gordon-Schneider for assistance in cell sorting and flow cytometry analysis, and the JP Sulzberger Columbia Genome Center for NGS support. We thank C. Lu and V. Sahu for cell sonicator instrument access and valuable discussions on ChIP-seq analysis. G.D.L. is supported by the Columbia Integrated in CMBS Training Grant T32GM008224-34. R.T.K. is supported by the Columbia University Medical Scientist Training Program grant T32GM007367. A.C. is supported by a Career Awards for Medical Scientists from the Burroughs Wellcome Fund and NIH grant HG011855-01 from NHGRI. S.H.S. is supported by NIH grant DP2HG011650-01, a Pew Biomedical Scholarship and Sloan Research Fellowship, and a generous start-up package from the Columbia University Irving Medical Center Dean’s Office and the Vagelos Precision Medicine Fund.

## AUTHOR CONTRIBUTIONS

G.D.L. performed western blots, plasmid-based reporter assays, and experiments screening CAST homologs and host proteins. R.T.K. performed genomic activation assays, ChIP experiments, and ddPCR analyses. G.D.L. and R.T.K. performed integration assays and quantified integration efficiency by qPCR and amplicon sequencing. T.S.H-H. performed initial human cell experiments and, together with M.I.H., tested tagged constructs for activity in *E. coli*. S.E.K. tested tagged constructs in *E. coli* and assisted with initial NGS library preparation and data analysis. P.L.H.V. designed and cloned constructs for initial I-E activation assays. S.T. performed ChIP-seq analysis. A.C. provided expert support and helped S.H.S. supervised the project. S.H.S., G.D.L., R.T.K., and A.C. discussed the data and wrote the manuscript, with input from all the authors.

## SUPPLEMENTARY FIGURES

**Supplementary Fig. 1.**
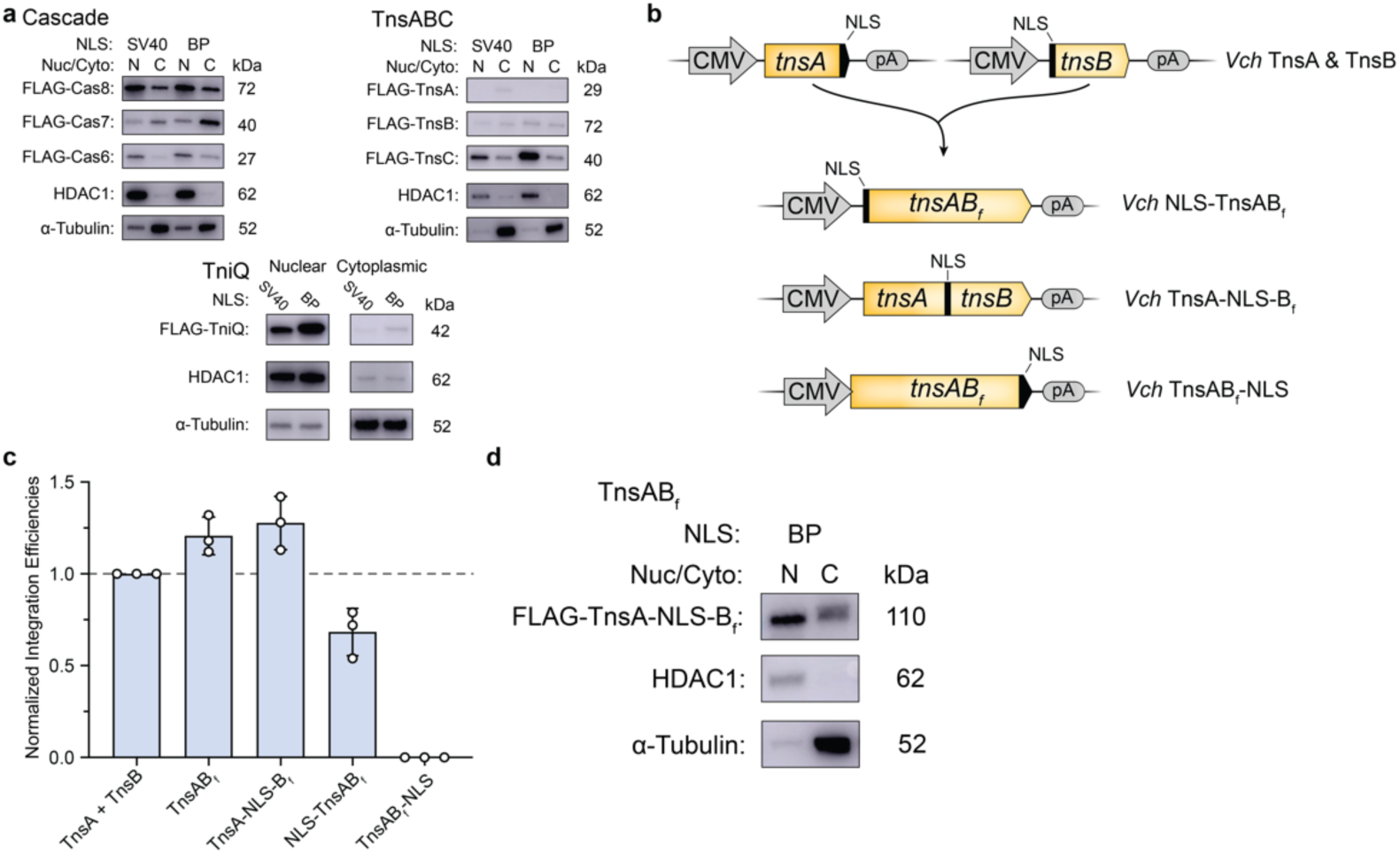
Improving expression and nuclear localization of *Vch*CAST components. **a,** Western blotting of various *Vch*CAST components using distinct nuclear localization signals (NLS). Each component was appended with a 3xFLAG epitope tag and NLS tag, and nuclear fractionation was performed to separate nuclear and cytoplasmic cellular proteins. Histone deacetylase 1 (HDAC1) and ɑ-Tubulin were used as nuclear- and cytoplasmic-specific loading controls, respectively. **b,** Multiple fusion designs of TnsA and TnsB (TnsAB_f_) were generated, with an NLS appended internally or at the N- or C-terminus. **c,** RNA-guided DNA integration activity was determined in *E. coli* with the indicated TnsABf variants, as measured by qPCR. **d,** Western blotting of TnsAB_f_ with internal NLS validated expression and nuclear localization. The observed band was at the expected size, with no evidence of degradation or internal cleavage.

**Supplementary Fig. 2.**
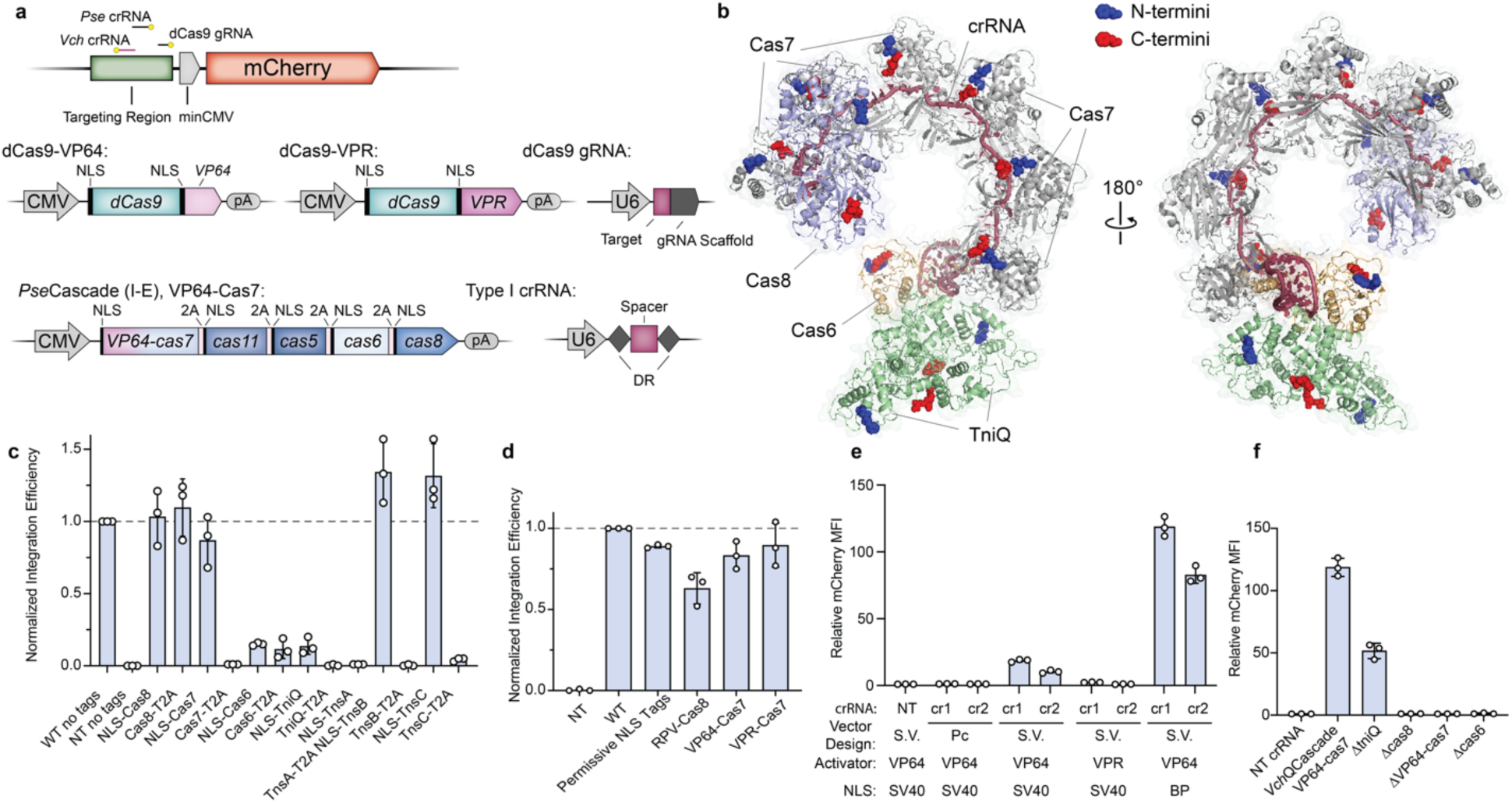
Optimization of *Vch*QCascade expression and transcriptional activation in human cells. **a,** Above, schematic of mCherry reporter plasmid for transcriptional activation assays. The location of sites targeted by Cas9 single-guide RNAs (sgRNA) and Cascade CRISPR RNAs (crRNA) are indicated. PAMs are marked with a yellow circle. Below, design of mammalian expression vectors encoding Cascade-based transcriptional activators from a Type I-E system (*Pse*Cascade)^40^, alongside dCas9-VP64 and dCas9-VPR controls^55, 61^ b, Depiction of *V. cholerae* TniQ-Cascade structure (PDB ID: 6PIF) showing the location of N- and C-termini in blue and red, respectively. All termini are solvent exposed and appear amenable to tagging. **c,** RNA-guided DNA integration activity in *E. coli* with the indicated NLS and/or 2A-tagged protein variants, measured by qPCR. Numerous tags have a deleterious effect. Data are normalized to the “WT no tags” condition, which resulted in a mean integration efficiency of 51 ± 8 %. **d,** RNA-guided DNA integration activity in *E. coli* with combined NLS and transcriptional activator fusions, as measured by qPCR. Fusing a VP64 or VPR transcriptional activator to the N-terminus of Cas7 exhibited the least deleterious effects on integration activity. Data are normalized to the “WT” condition, which resulted in a mean integration efficiency of 76.4% ± 4%. **e,** Strength of transcriptional activation across a set of distinct crRNAs (“cr#”) targeting the mCherry reporter plasmid, as well as various activator-NLS constructs. Activation was measured using the reporter shown in **Supplementary Figure 2a** and measured by flow cytometry. S.V. indicates single vector design. Pc indicates polycistronic design of expression vectors as shown in **Supplementary Figure 2a f,** Transcriptional activation by *Vch*QCascade utilizing a VP64-Cas7 fusion construct is strictly dependent on the presence of all Cascade components, as seen from the indicated dropout panel, but proceeds with ∼50% activity in the absence of TniQ. Data in **c–f** are shown as mean ± s.d. for n = 3 biologically independent samples.

**Supplementary Fig. 3.**
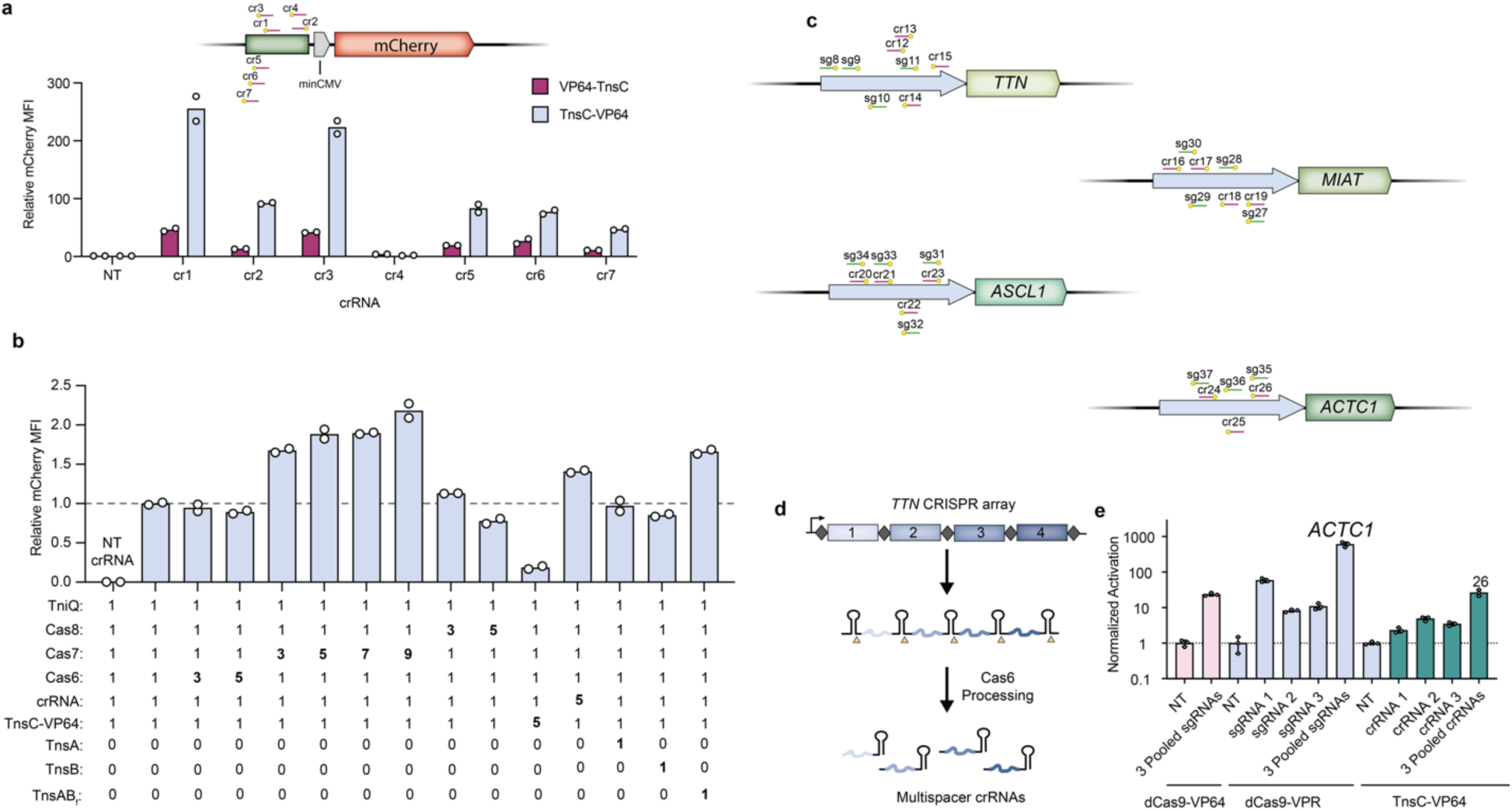
Optimization of TnsC-mediated transcriptional activation in human cells. **a,** Normalized mCherry fluorescence levels for the indicated experimental conditions, as measured by flow cytometry. VP64 was appended to TnsC at either the N- or C-terminus (VP64- TnsC or TnsC-VP64, respectively), and crRNAs (“cr#”) were cloned to target various sites upstream of the mCherry gene (top). mCherry fluorescence levels were measured by flow cytometry and normalized to the non-targeting gRNA condition (bottom). **b,** Transcriptional activation is affected by titrating the relative levels of each expression plasmid, with numbers below the graph indicating the fold-change of each plasmid amount relative to the initial stoichiometric condition with a targeting crRNA (second bar from left). mCherry fluorescence levels were measured by flow cytometry. **c,** Schematic showing the position of crRNAs (“cr#”) or sgRNAs (sg#) targeting each genomic locus for TnsC-mediated transcriptional activation for *Vch*CAST (maroon) and dCas9 *TTN* activation (green). **d,** Representative schematic of multispacer crRNAs used during TnsC-mediated genomic transcriptional activation. For *TTN*, *MIAT*, and *ASCL1*, the 4 individual spacer sequences used in individual or pooled crRNA conditions were expressed as one multispacer CRISPR array. The CRISPR array is processed by Cas6 after transfection into cells and enables programmable targeting of multiple copies of QCascade and TnsC to a target locus **e,** Genomic transcriptional activation at the *ACTC1* locus as quantified by RT-qPCR. 3 distinct crRNAs or gRNAs were used for each condition. Data in **a**,**b** are shown as mean for n = 2 biologically independent samples. Data in **e,** are shown as the mean ± s.d. for n = 3 biologically independent samples.

**Supplementary Fig. 4.**
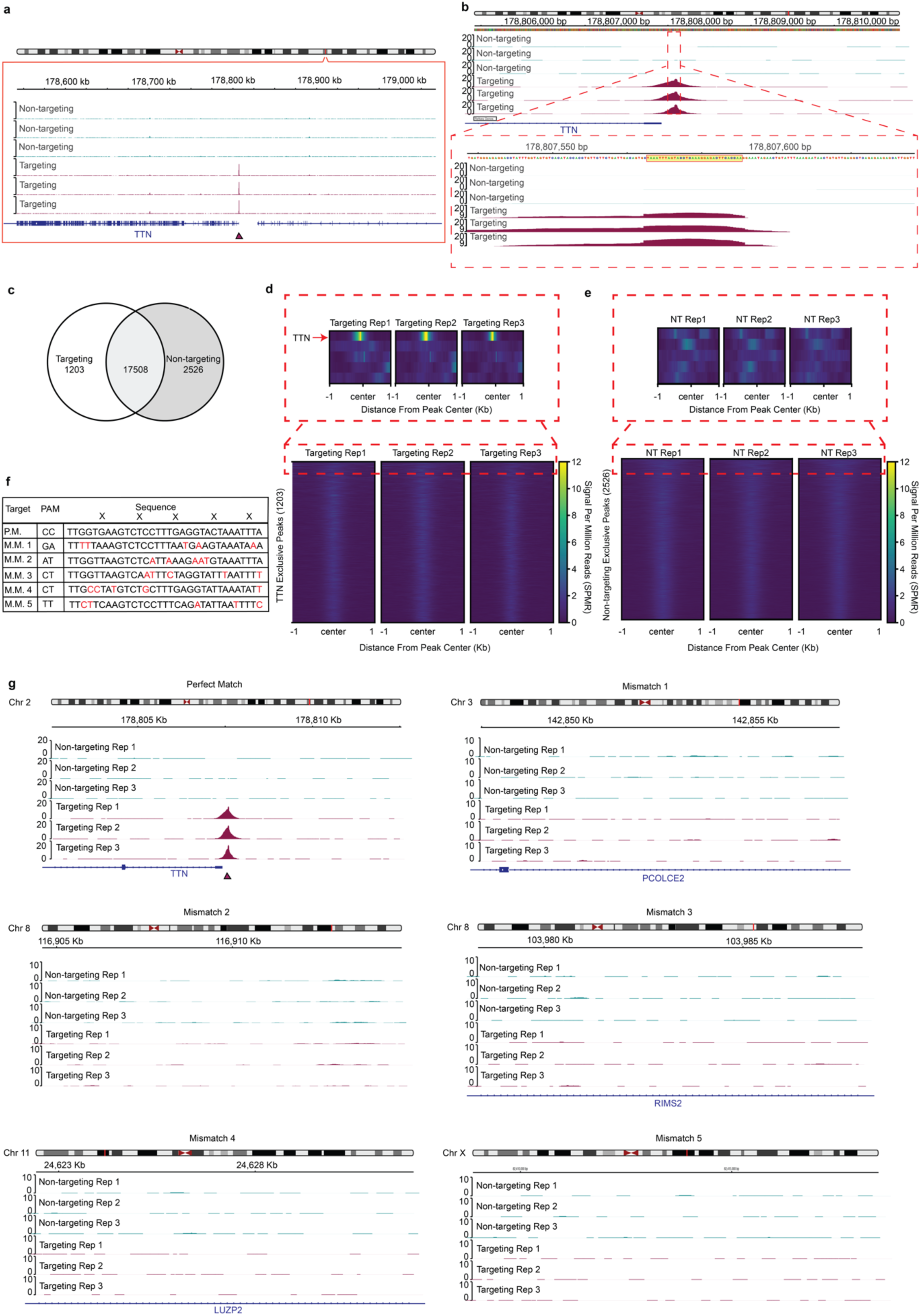
Detection of TnsC recruitment to a genomic locus and profiling of off-target binding events. **a,** 500 kb viewing window of ChIP-seq signal at the *TTN* promoter targeted by *TTN* Guide 1. **b,** Top, 5 kb viewing window of ChIP-seq peak at the *TTN* promoter targeted by TTN Guide 1. Bottom, 150 bp viewing window ChIP-seq peak at the *TTN* promoter targeted by TTN Guide 1. The peak summits in the targeting conditions align with the *TTN* promoter protospacer. **c,** Venn diagram showing overlap of targeting and non-targeting peaks. **d,** Heatmap of signal intensity in a 2 kb window surrounding the peak center in *TTN* targeting exclusive peaks, sorted in descending order by mean signal over the window. The peak with the highest mean signal was at the *TTN* promoter, which was targeted by *TTN* Guide 1. **e,** Heatmap of signal intensity in a 2 kb window surrounding the peak center in non-targeting (NT) exclusive peaks, sorted in descending order by mean signal over the window. ChIP-seq signal was weak across NT exclusive peaks. **f,** List of 5 genomic loci most similar to the *TTN* protospacer. Mismatches at every 6th nucleotide, denoted by an “X”, were disregarded due to the nature with which Cas7 binds to crRNAs. All other mismatches are shown in red. **g,** Manual inspection of a 10 kb window surrounding each predicted mismatch sequence. Minimal enrichment of ChIP-seq signal was seen in either the *TTN* targeting or the non-targeting condition. Triangles in **a**, **g**, denote the position of either the expected *TTN* targeting sequence or of the predicted mismatch sequences.

**Supplementary Fig. 5.**
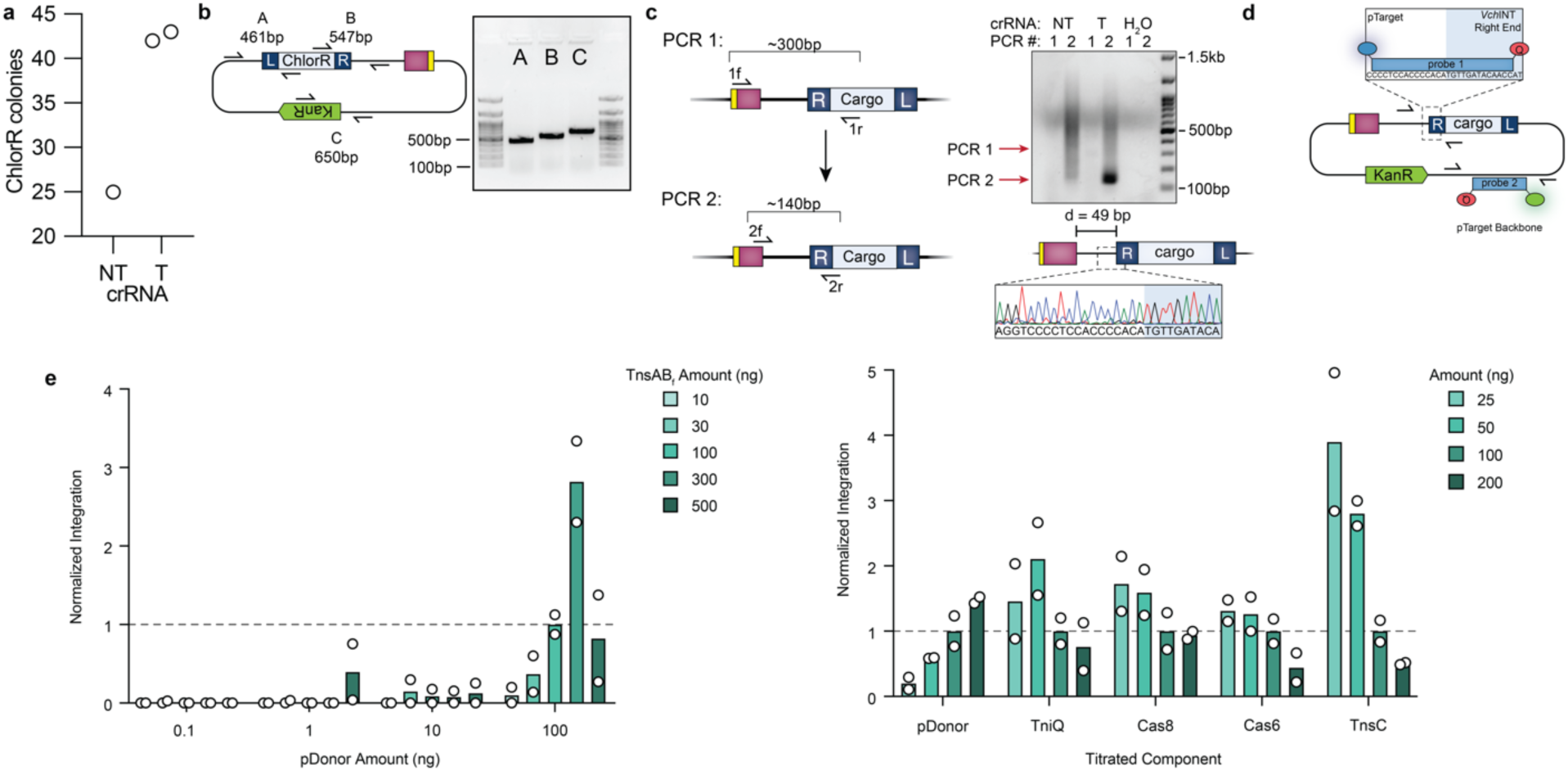
Initial detection and optimization of targeted integration using *Vch*CAST (eCAST-1). **a,** Initial quantification of ChlorR resistant *E. coli* colonies after isolation from human cells. **b,** Representative colony PCR of clonal transposition products, detecting TnR and TnL junctions, as well as the KanR marker on the backbone of pTarget. Sanger sequencing of integration junctions are shown in Figure 4b. **c,** Nested PCR strategy to detect plasmid-transposon junctions directly from HEK293T cell lysates (left), and agarose gel electrophoresis showing target-cargo junction product bands (right). Expected amplicon sizes are marked for each PCR reaction with red arrows, and the crRNA was either non-targeting (NT) or targeting (T). “H_2_O” denotes a condition in which the lysate was omitted from the PCR reactions. An aliquot of PCR is used for PCR 2 such that a “nested PCR” is performed (see Methods). Sanger sequencing was performed on the product after PCR 2 in the targeting condition (bottom right). **d,** Schematic of TaqMan probe strategy used to improve signal-to-noise by selectively detecting novel plasmid-transposon junctions. Probes labeled with FAM (blue) are used to detect target-transposon junctions, and probes labeled with SUN (green) are used to detect the target plasmid backbone, for integration efficiency quantification. Probes that span the junction of pTarget and the right transposon end of *Vch*CAST are designed to anneal to an insertion event 49-bp downstream of the target site. **e,** Integration efficiencies were improved by varying the relative levels of pDonor, pTarget, or protein expression plasmids, as indicated; data were normalized to a control sample transfected with 100 ng of each component (left), or to the 100 ng condition for each varied protein (right), which had an average value of either 0.004 % (left) or ranged from 0.0002–0.0005 % (right), respectively. Data in **e** are shown as mean for n = 2 biologically independent samples. Data shown in **e** are quantified via qPCR.

**Supplementary Fig. 6.**
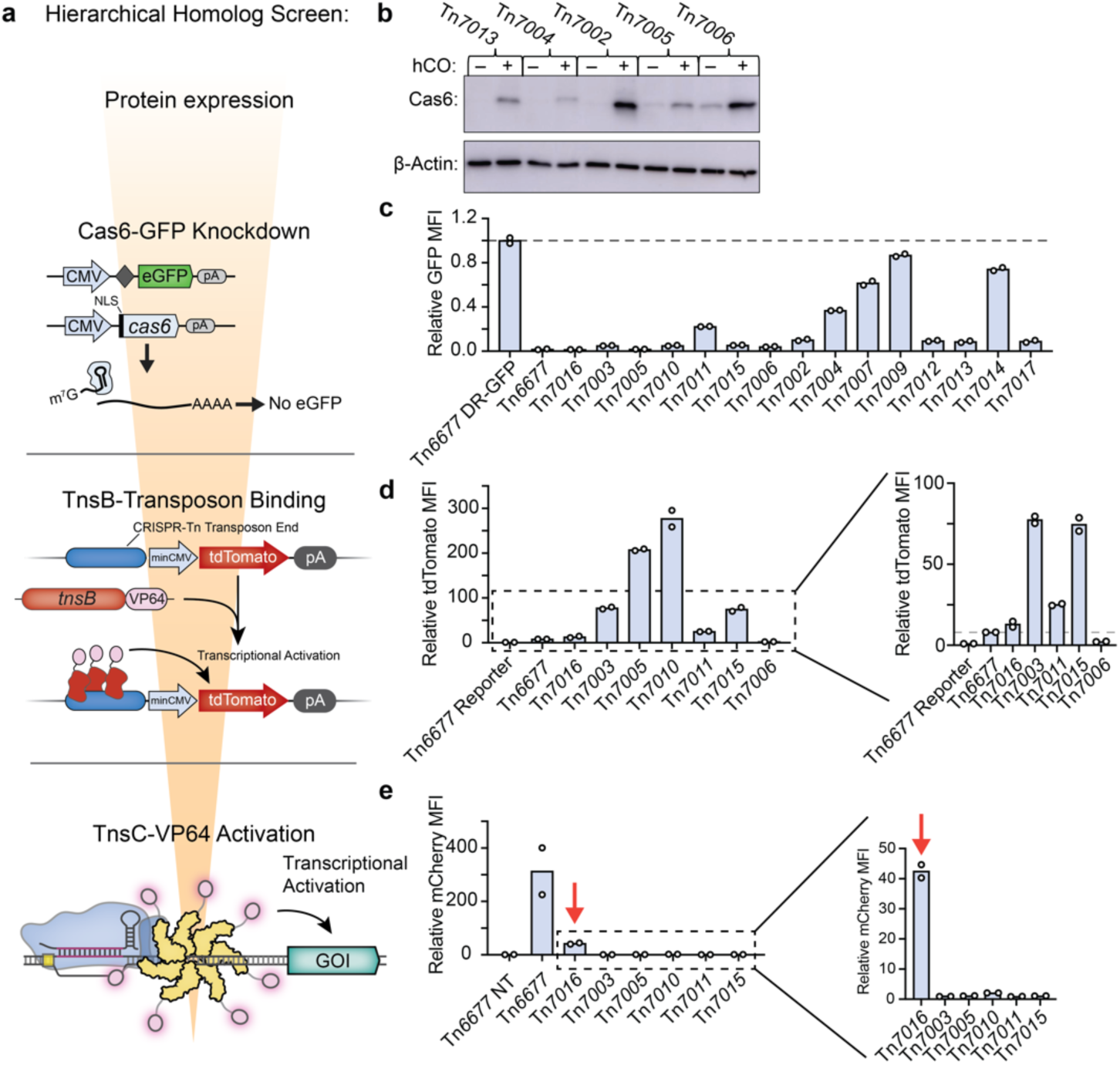
Systematic screening of homologous Type I-F CRISPR-associated transposons to uncover improved systems for mammalian cell applications. **a,** Cartoon depicting the multi-tiered approach that was applied to screen the indicated systems through a series of consecutive activity assays, with associated schematics shown for each functional assay. The middle panel depicts a transcriptional activation assay designed to monitor transposon DNA binding by TnsB in human cells using a tdTomato reporter plasmid. **b,** Western blotting to detect expression of candidate Cas6 homologs in HEK293T cells, with or without human codon optimization (hCO), using anti-FLAG antibody; β-actin was used as a loading control. We observed a range of expression levels for human codon-optimized gene variants, and genes were poorly expressed for most systems when native bacterial coding sequences were used. **c,** Activity assays for Cas6 homologs using the GFP knockdown assay shown in Figure 1d. For each homolog, GFP fluorescence levels were measured by flow cytometry and normalized to the experimental condition in which the GFP reporter plasmid lacked a CRISPR direct repeat (DR) in the 5’-UTR. **d,** Transcriptional activation data for TnsB-VP64 constructs from selected homologous CRISPR-associated transposons, as measured by flow cytometry. **e,** Transcriptional activation data for QCascade and TnsC-VP64 from homologous CRISPR-associated transposons, as measured by flow cytometry. Tn*7016*, the final homolog that was selected for additional screening for transposition, is marked with a red arrow. Data in **c–e** are shown as mean for n = 2 biologically independent samples.

**Supplementary Fig. 7.**
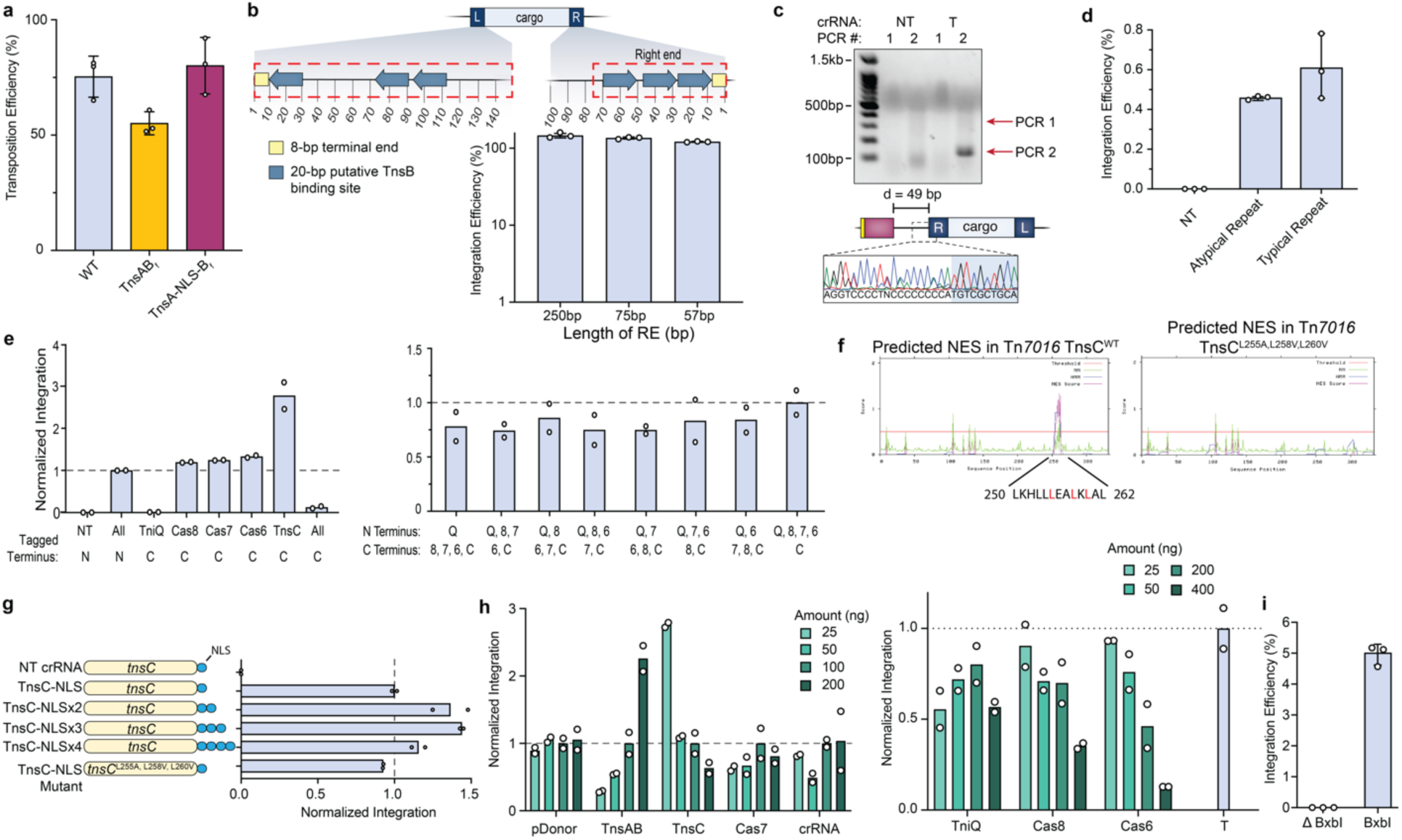
Parameter screening to further improve integration activity with the eCAST-2 (*Pse*CAST) system. **a,** RNA-guided DNA integration efficiency for TnsAB fusion (TnsAB_f_) protein design, with or without internal NLS, compared to the wild-type TnsA and TnsB proteins. Experiments were performed in *E. coli*, and efficiencies were measured by qPCR. **b,** Tn*7016* transposon ends were shortened relative to the constructs tested previously^36^, generating the constructs indicated with red dashed boxes at the top. RNA-guided DNA integration activity was compared for the indicated transposon right end (RE) variants in *E. coli*, as measured by qPCR (bottom), while a 145bp LE was used. The final pDonor design used in Figure 4 contains 145-bp and 75-bp derived from the native left and right ends of *Pseudoalteromonas* Tn*7016*, respectively. **c,** Agarose gel electrophoresis showing successful junction products from nested PCR (top) for *Pse*CAST, and Sanger sequencing chromatograms showing the expected integration distance (bottom). **d,** Integration efficiencies in HEK293T cells were similar using either typical or atypical CRISPR repeats^39, 67^, as measured by qPCR. **e,** RNA-guided DNA integration activity was compared with the indicated BP NLS tags on *Pse*CAST components, as measured by qPCR. Individual components had their respective BP NLS tag repositioned from the N- to the C-terminus; “All” represents a condition in which all components had BP NLS tags on the noted terminus (left). Interestingly, the observed tag sensitivity is similar to, but distinct from, that with *Vch*CAST components. Various combinations of N- and C-terminal NLS tagging for *Pse*QCascade and *Pse*TnsC (right). NT = non-targeting crRNA. **f,** Nuclear export signal (NES) predictions for *Pse*CAST wild type (WT) and mutant TnsC. We hypothesized that a putative NES within TnsC could led to inefficient nuclear localization, and selected multiple residues that, when mutated, might lower this risk. Predicted NES sequences were generated using NetNES. **g,** RNA-guided DNA integration activity was compared after appending additional NLS tags on *Pse*TnsC and removing a potential internal nuclear export signal (NES) sequence, as indicated in panel **f**. **h,** RNA-guided DNA integration activity was compared after varying the relative levels of individual *Pse*CAST protein and RNA expression plasmids. Data were measured by qPCR and are normalized to either the sample transfected with 100 ng of each component for each condition, with an average integration efficiency of 0.10–0.17% (left), or a control sample (labeled T) transfected with the standard *Pse*CAST plasmid amounts, as detailed in the Methods section with an average integration efficiency of 2.7 % (right). **i,** A plasmid-based BxbI recombination assay was performed to benchmark *Pse*CAST integration efficiency to other commonly used large DNA insertion tools. Data in **a,b**,**d, i** are shown as the mean ± s.d. for n = 3 biologically independent samples. Data in **e**,**g**,**h** are shown as the mean for n = 2 biologically independent samples.

**Supplementary Fig. 8.**
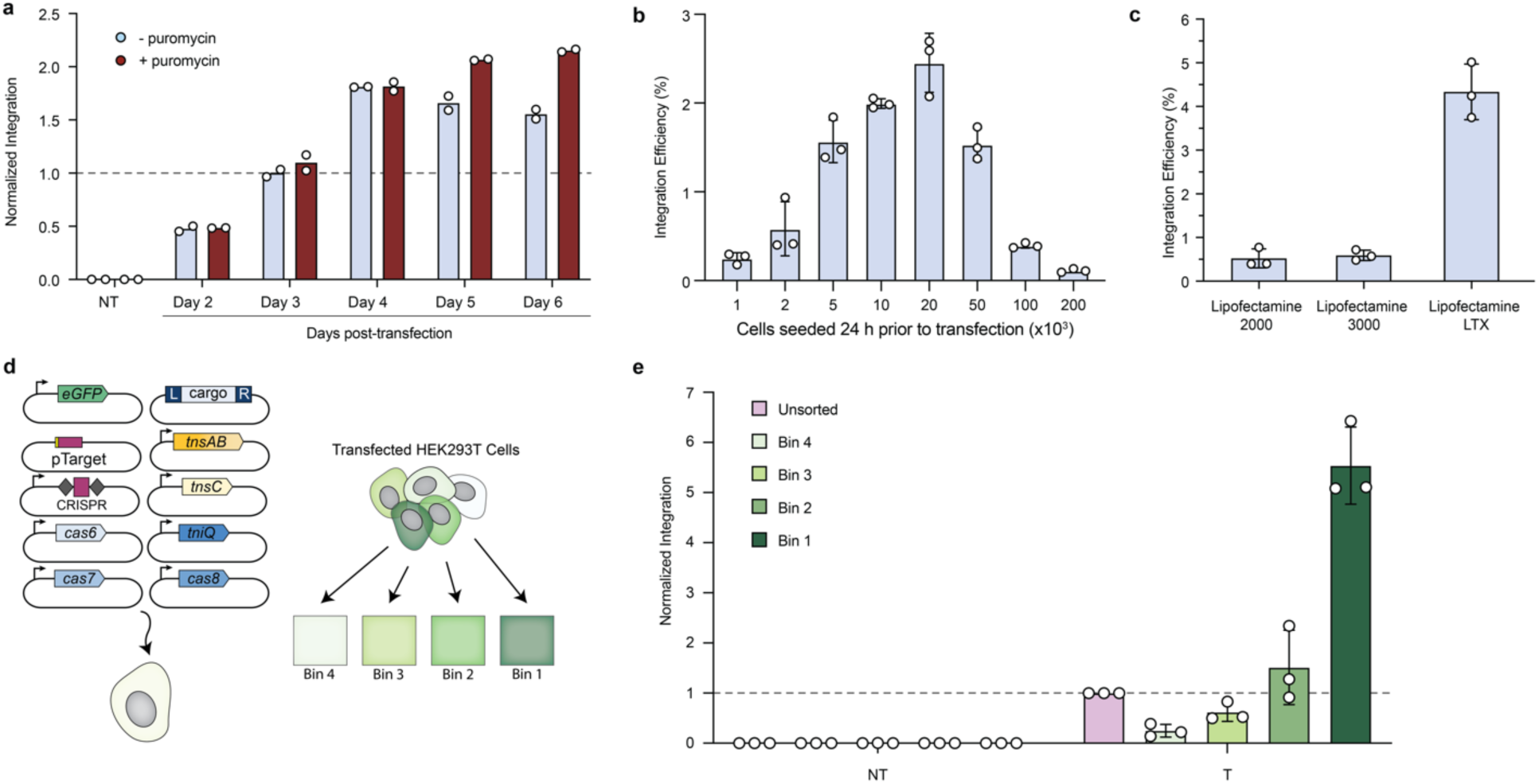
Selection, seeding, and sorting strategies result in further increases in eCAST-2.2 integration efficiencies. **a,** Normalized RNA-guided DNA integration efficiency for eCAST-2.2 in the absence or presence of puromycin selection, and after harvesting cells from between 2–6 days post-transfection. Experiments used a puromycin resistance plasmid as a transfection selection marker, in addition to eCAST-2.2 component plasmids, and integration activity was measured by qPCR and normalized to the condition harvested on day 3 without puromycin selection, which had an average integration efficiency of 2.3 %. **b**, eCAST-2.2 integration efficiencies were compared as a function of seeding density 24 hours before transfection. 24-well plates were with various cell densities ranging from 10^3^ to 2 x 10^5^ cells per well, and integration activity was measured by qPCR. **c**, Transfection of HEK293T cells via various cationic lipid delivery methods affected integration efficiencies. **d**, Schematic showing the use of a GFP transfection marker and cell sorting to increase integration efficiency. A GFP expression plasmid was transfected in significantly smaller amounts relative to eCAST-2.2 component plasmids, and cells were sorted into bins of varying GFP expression levels. **e**, eCAST-2.2 integration efficiencies are enhanced after using flow cytometry to sort cells for the brightest GFP positive cells. Cells were sorted four days after transfection, and the top 20% brightest cells were binned in increments of 5%, with Bin 1 representing the top 5% brightest cells and Bin 4 representing the 15-20% brightest cells. Integration efficiencies were determined for each bin separately, or for the unsorted population, as measured by qPCR. Integration efficiencies were normalized to the unsorted, targeting crRNA condition, which had a value of 0.44 %. Data in **a** are shown as the mean of n = 2 biologically independent samples. Data in **b**, **c**, **e** are shown as the mean ± s.d. for n = 3 biologically independent samples.

**Supplementary Fig. 9.**
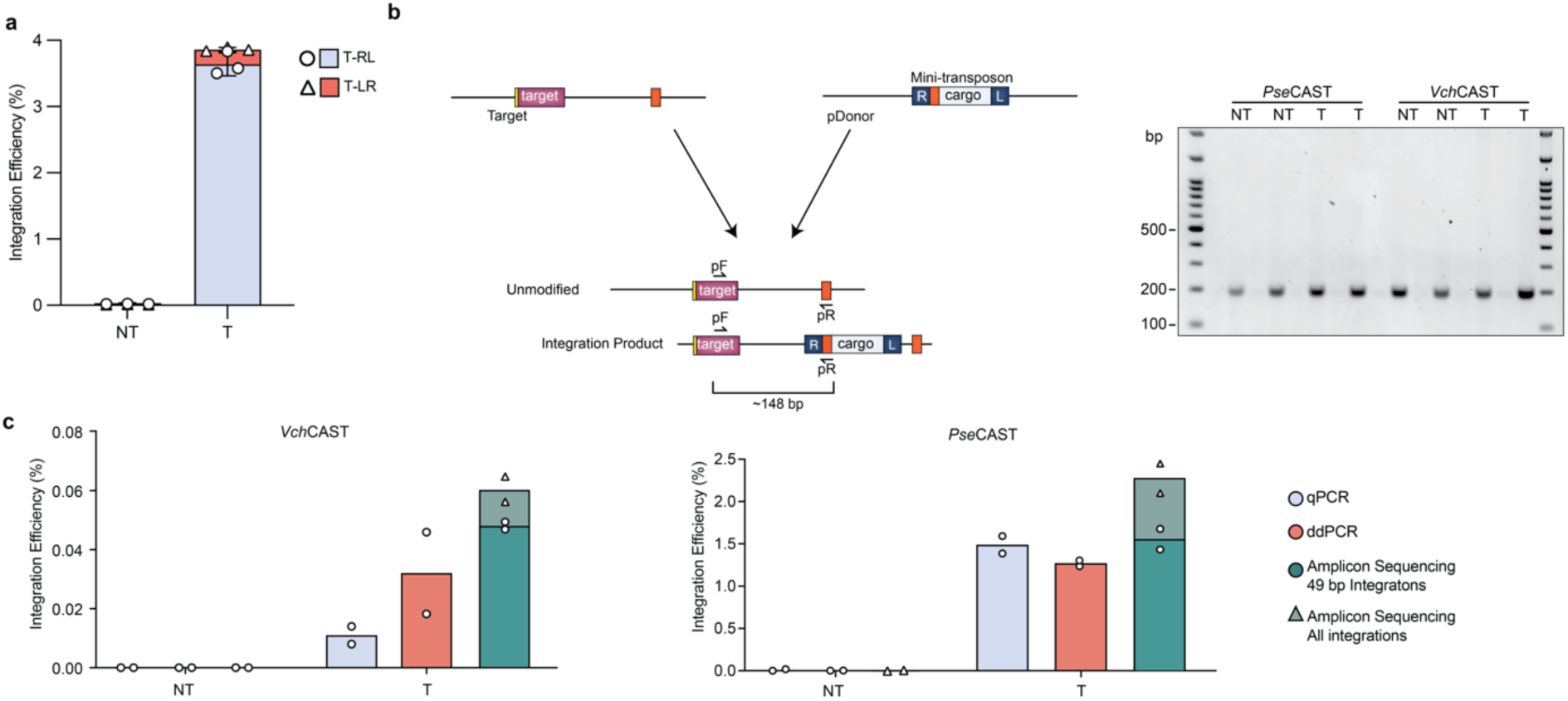
eCAST-2.2 integration is biased towards tRL insertion and reproducibly quantified across distinct approaches. **a**, RNA-guided DNA integration is heavily biased towards insertion in the right-left (tRL) orientation, with only a small minority of insertion events occurring in the left-right (tLR) orientation. Integration efficiencies were calculated using SYBR qPCR. **b**, Strategy to detect and quantify integration efficiencies using PCR and next-generation sequencing. A variant pDonor was construct, in which a primer binding site is present within the transposon cargo at a distance from the transposon right end (R), such that unintegrated and integrated pTarget molecules yield amplicons of indistinguishable length using pF and pR primers (left). Consequently, next-generation sequencing of these amplicons can provide relative ‘counts’ of edited and unedited alleles in the population, without introduction of PCR bias. Agarose gel electrophoresis demonstrates identical amplicon products for non-targeting (NT) and targeting (T) samples after PCR 1 for NGS analysis (right). **c**, Calculated integration efficiencies for the same experimental samples, measured by TaqMan qPCR, droplet digital PCR (ddPCR), and amplicon deep sequencing. ddPCR and qPCR analyses specifically probe for integration products that are 49-bp downstream of the target site, whereas amplicon sequencing analysis does not impose the same stringent distance bias, allowing the quantification of integration products within a larger window surrounding the anticipated integration site. Editing efficiencies for both eCAST-2.2 and eCAST-1 were consistent between different quantification methods. Data in **a**, are shown as the mean ± s.d. for n = 3 biologically independent samples. Data in **c** are shown as the mean for n = 2 biologically independent samples.

**Supplementary Fig. 10.**
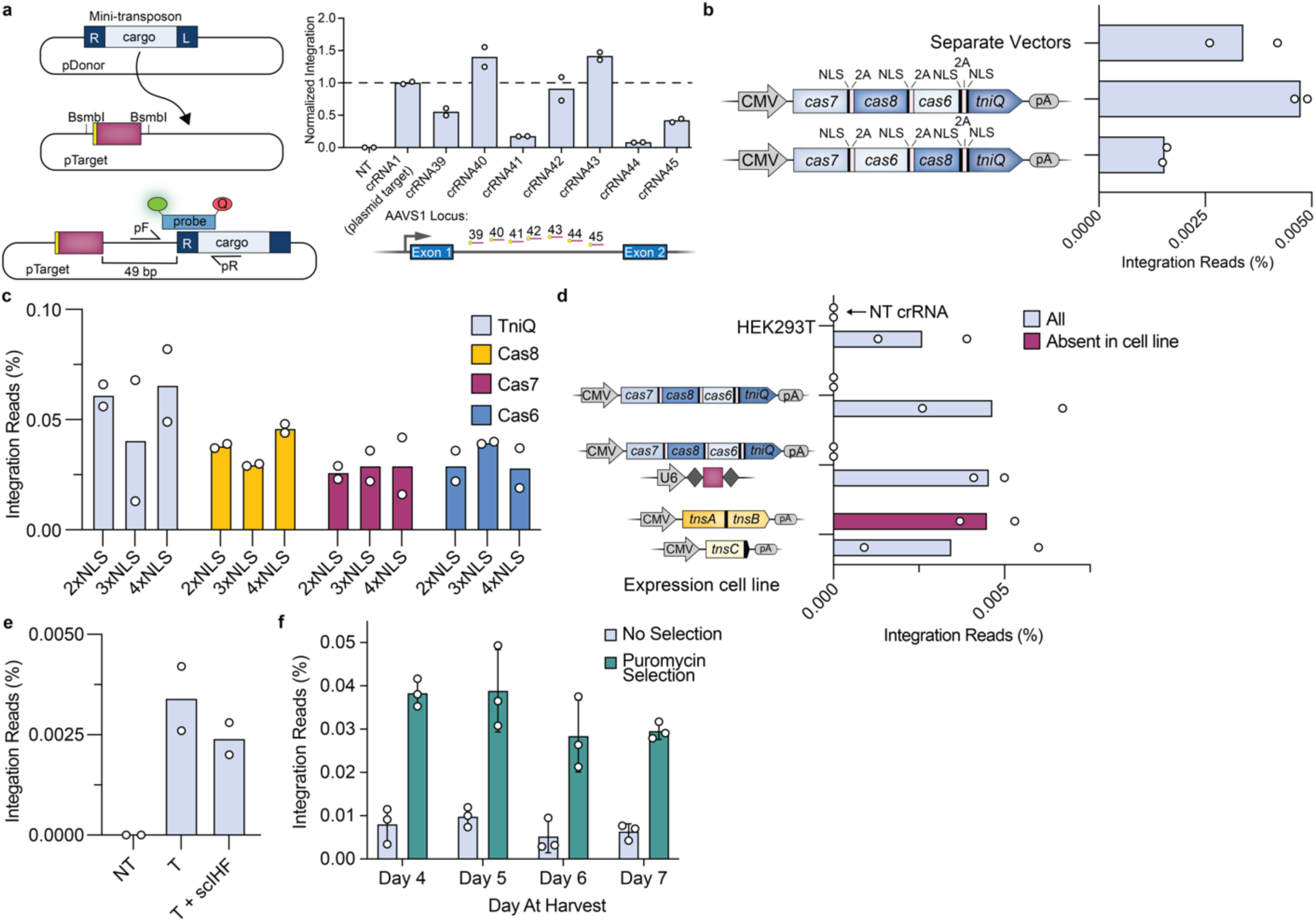
Preliminary attempts to improve eCAST-2.2 genomic integration activity and identify kinetic bottlenecks. **a,** A unique target site was cloned into a modified pTarget, in which the downstream integration site sequence remained the same, allowing us to specifically understand the impact of different crRNA sequences on integration efficiencies (left). Cloning various target sites into our modified pTarget that correspond to target sites within the AAVS1 safe harbor locus enabled screening of crRNAs to identify active sequences (right). Efficiencies were normalized to the crRNA used in our plasmid-targeting assays, which had an average integration efficiency of 2.0 %. **b**, Simplification of transfection workflow via polycistronic expression of QCascade, and genomic integration efficiencies with different constructs. “Separate Vectors” represents a condition in which *TniQ, Cas8, Cas7,* and *Cas6* were all expressed from separate pcDNA3.1-like vectors. **c,** Impact of additional NLS tags on *Pse*CAST QCascade components on genomic integration efficiencies. All QCascade components had a singular NLS tag, unless noted. **d,** Impact of stably-expressed *Pse*CAST components on genomic integration efficiencies. Cell lines were generated via Sleeping beauty with drug selection, and various components were stably expressed (indicated by operons shown on the y-axis). “All” represents conditions in which all *Pse*CAST components were co-transfected, while “Absent in cell line” represents conditions in which only the non-expressed *Pse*CAST components were transfected. **e,** Impact of cotransfection of *E. coli* derived Integration Host Factor (IHF) on human genomic integration efficiencies. “scIHF” represents a condition in which a plasmid expressing a single-chain IHFa/b^93^ was co-transfected. **f,** Varying cell harvest day and selection of transfected cells based on a concurrent drug marker improves integration efficiencies, although overall efficiencies remain low. Data in **a,-f,** are shown as mean for n = 2 biologically independent samples. Data in **g,** are shown as the mean ± s.d. for n = 3 biologically independent samples. Data in **a,** and ChIP-qPCR data in **b,** were determined by qPCR. Data in **b, d – g,** were determined by amplicon sequencing.

**Supplementary Fig. 11.**
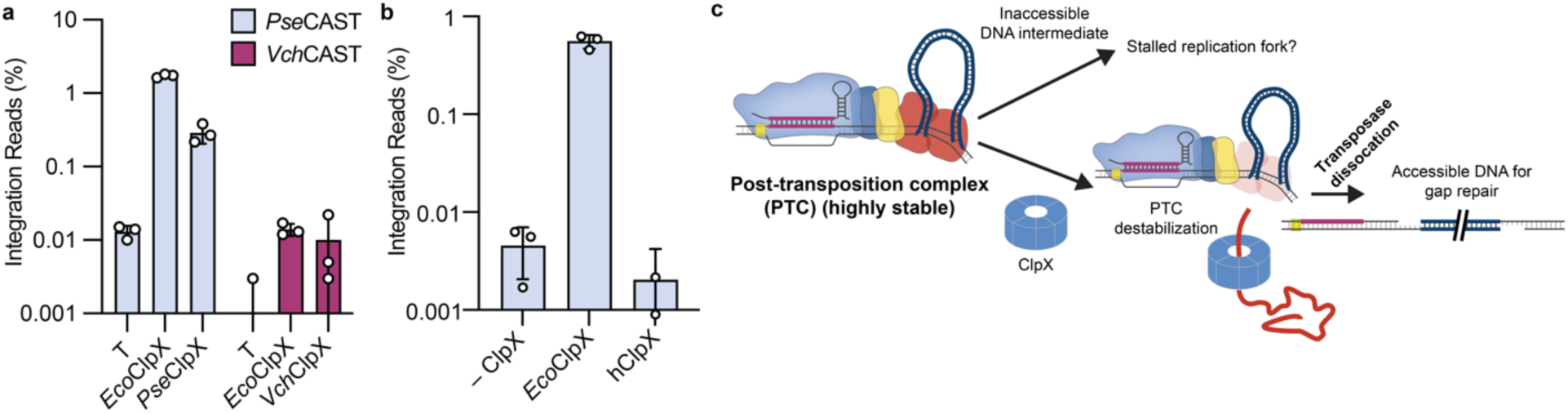
Genomic editing outcomes with ClpX. **a,** The impact of native ClpX proteins on *Pse*CAST and *Vch*CAST. *Pse*ClpX and *Vch*ClpX improved *Pse*CAST and *Vch*CAST genomic integration efficiencies, respectively, but *Eco*ClpX consistently produces a more robust improvement. **b,** Human-derived ClpX does not improve genomic integration efficiencies for *Pse*CAST. The putative mitochondrial targeting sequence from human derived ClpX^94, 95^ was replaced with a BP-NLS tag. **c,** Proposed model for ClpX’s role in improving genomic integration efficiencies. In the absence of ClpX, the PTC is sufficiently stable to prevent accessibility to the DNA intermediate, leading to a loss of genomic integration events. In contrast, including ClpX unfolds CAST components, resulting in destabilization of the complex and accessibility to the DNA intermediate. Data in **a, b** are shown as the mean ± s.d. for n = 3 biologically independent samples.

**Supplementary Fig. 12.**
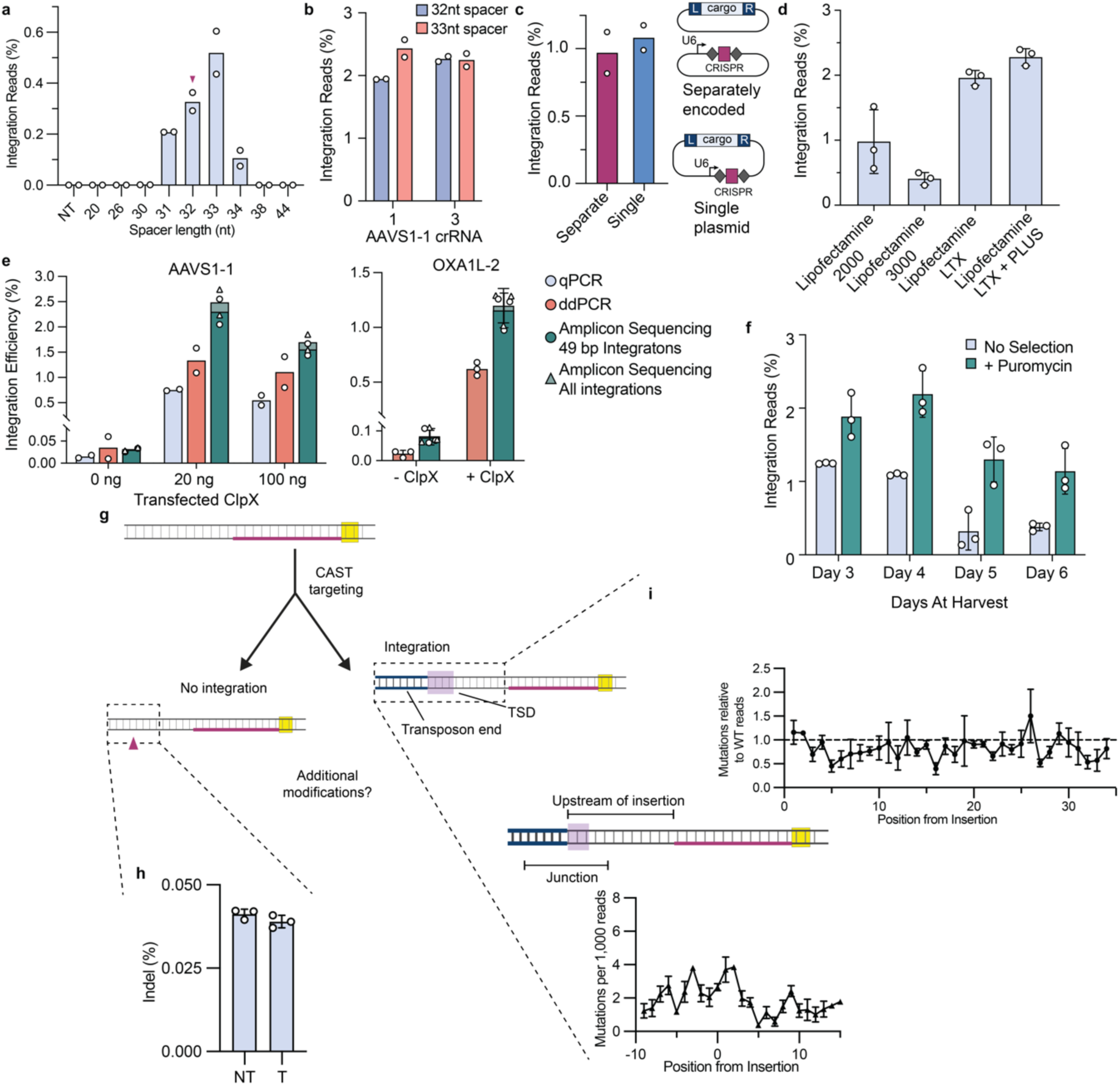
Engineering CAST systems with ClpX. **a**, Impact of atypical spacer lengths on plasmid-based integration efficiencies (the canonical spacer length, 32nt, is marked with a maroon triangle). **b,** Impact of 32nt vs 33nt spacer lengths on genomic integration efficiencies at the AAVS1-1 target site. Two different crRNAs were tested that were nearby in the genomic locus, minimizing disruption of potential downstream integration-site requirements. **c,** Impact of encoding the crRNA on the pDonor for genomic integration efficiencies. The U6 promoter, crRNA, and U6 terminator sequences were cloned on either a separate plasmid or in the pDonor backbone. **d,** Genomic integration as a function of different cationic lipid transfection methods **e,** A comparison of integration efficiencies in the presence and absence of ClpX as measured by qPCR, ddPCR, and amplicon sequencing for AAVS1-1; ddPCR and amplicon sequencing for OXA1L-2. **f,** Varying cell harvest day and selection of transfected cells based on a concurrent drug marker improves integration efficiencies, in the presence of ClpX **g,** Schematic of sequences that were analyzed to understand if undesirable editing outcomes were occurring with eCAST-3. If a sequence did not contain a transposon end, the sequence surrounding the intended integration site was investigated for a higher frequency of indel events compared to samples in which a non-targeting crRNA was used. If a transposon end was detected in the sequence, the sequence was analyzed for additional mutations. **h,** Mutations surrounding the integration region do not occur above background frequencies present when a NT crRNA is co-transfected. **i,** Mutations upstream the integration site do not occur at a higher rate compared to WT alleles (top). Mutations in the transposon end and surrounding the target site duplication do not occur at rates above background sequencing error (bottom). Data in **a-c, e (AAVS1-1)** are shown as mean for n = 2 biologically independent samples. Data in **d, e (OXA1L-2), f, h, i** are shown as mean ± s.d. for n = 3 biologically independent samples. Data are quantified by amplicon sequencing.

**Supplementary Fig. 13.**
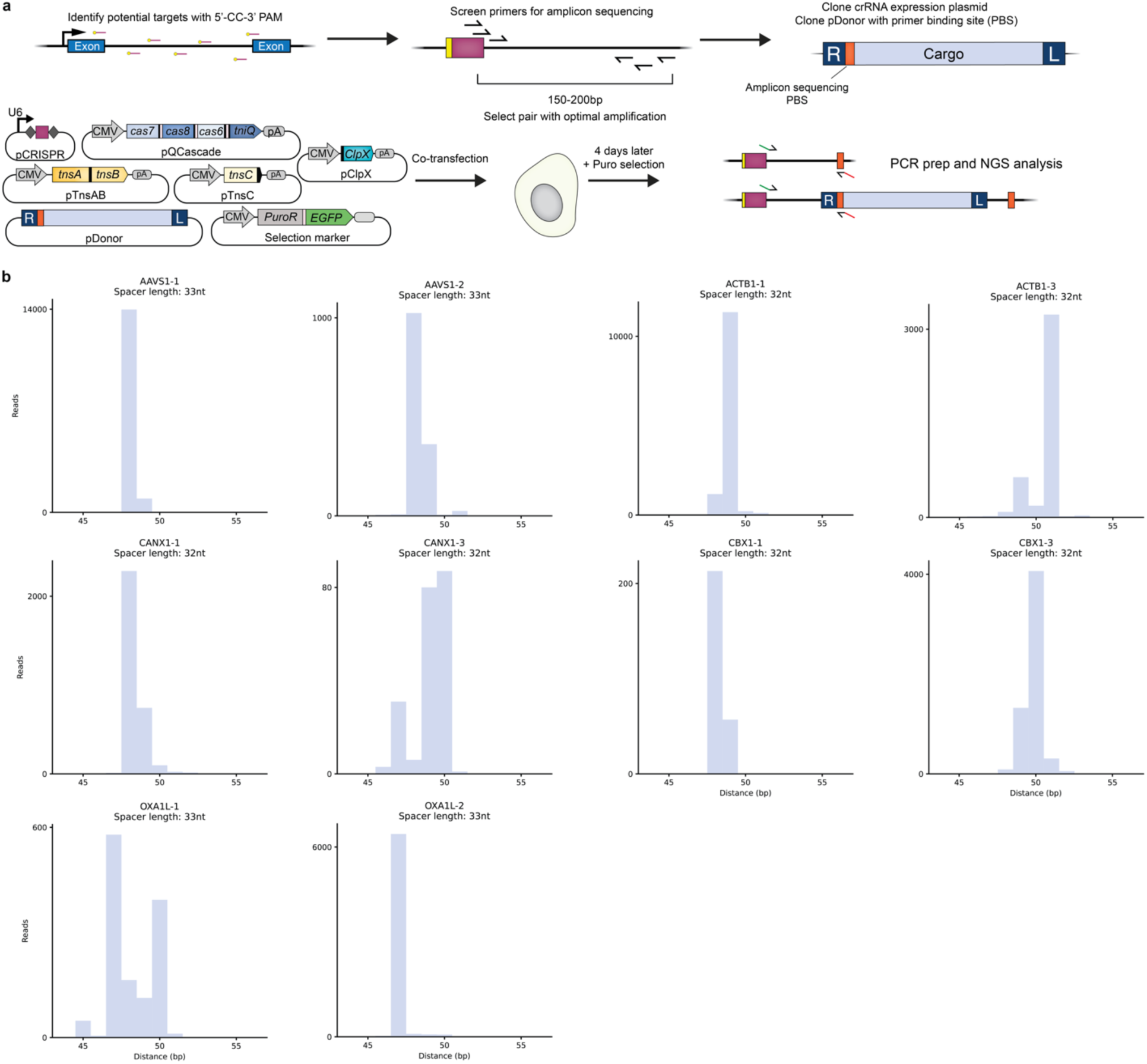
Leveraging eCAST-3 to perform targeted RNA-guided DNA integration at multiple target sites. **a,** Current workflow for applying eCAST-3 to new target sites. First, potential targets with CC PAMs are identified in region of interest. Target sites are then screened for optimal primers for amplicon sequencing. The downstream primer binding site is cloned into a pDonor immediately adjacent to the RE, enabling NGS-based quantification. Cells are then transfected with pCRISPR, pQCascade, pTnsAB, pTnsC, pClpX, pDonor, and an optional drug selection marker. After 4 days, cells can be harvested for PCR prep and subsequent NGS-based analysis. **b,** Representative integration site distributions for transfections shown in Fig. 5i. The length of the spacer is shown, and the distance represents the length from the PAM-distal end of the spacer to the transposon end.

